# A large inner membrane pore defines the ESX translocon

**DOI:** 10.1101/800169

**Authors:** Nicole Poweleit, Nadine Czudnochowski, Rachael Nakagawa, Kenan Murphy, Christopher Sassetti, Oren S. Rosenberg

## Abstract

The ESX (or Type VII) secretion systems are protein export systems in mycobacteria and many Gram-positive bacteria that mediate a broad range of functions including virulence, conjugation, and metabolic regulation. These systems translocate folded dimers of WXG100-superfamily protein substrates across the cytoplasmic membrane; however, the architecture and mechanism of translocation has remained elusive. We report the cryo-electron microscopy structure of an ESX-3 system, purified using an epitope tag inserted with recombineering into the model organism *Mycobacterium smegmatis*. The structure reveals two large *α*-helical membrane pores of sufficient diameter to secrete folded substrates. A complex, asymmetric, multimeric cytoplasmic domain is poised to gate and regulate the pore’s function. Our study provides mechanistic insights into the ESX systems and will guide structure-based design of drugs targeting this unique bacterial translocon.

**One Sentence Summary:** The structure of the ESX-3 secretion system reveals a pore of sufficient size for the transit of folded substrates and a complex, cytoplasmic regulatory apparatus.

## Main Text

Mycobacteria use a set of specialized secretion systems called ESX to transport proteins across their complex, diderm cell walls (*1*). Originally described as virulence factors in *Mycobacteria tuberculosis* (*2*–*5*), orthologs of ESX have since been discovered in most gram-positive bacteria (*6*), and are more generally referred to as Type VII secretion systems (*7*). In mycobacteria there are five paralogous ESX operons (ESX 1-5) each of which encodes an inner membrane translocon complex consisting of three conserved Ecc proteins: EccB, EccC, and EccD. A fourth protein, EccE is conserved in all ESX operons except the ancestral ESX-4 operon and is also considered a part of the ESX translocon complex as it copurifies with EccB, EccC, and EccD (*8*). All Type VII secretion systems translocate proteins in the WXG100-superfamily, which share a common two-helix hairpin structure and are found as homo- or heterodimers (*9*) and are mutually dependent for secretion with other substrates (*10*). In contrast to the general (Sec) secretory apparatus, ESX substrates have been shown to be secreted in their folded, dimeric state (*11*).

Structural and functional information has been reported for truncated and isolated, soluble domains of the ESX translocon complexes and their homologs (*12*–*15*). A low resolution, negative stain electron microscopy structure of ESX-5 shows the translocon complex assembles into a hexamer (*16*). Structures of other proteins encoded in ESX operons including secreted substrates (*17*), substrate chaperons (*18*), and the regulator protease MycP (*19*) have been solved. Despite revealing important functional information about ESX, structures of overexpressed and isolated proteins are insufficient to understand how the translocation pore mediates the regulated secretion of fully folded substrates. We therefore undertook structural studies of a native ESX-3 complex from the model organism *M. smegmatis* using cryo-electron microscopy (cryo-EM).

The ESX-3 translocon complex is important for iron acquisition (*20, 21*), cell survival (*22*), and virulence in pathogenic mycobacteria (*23*), and its role in iron homeostasis is conserved in the model system, *M. smegmatis*. (*20*) The ESX-3 translocon complex proteins are transcribed in a single operon (*24*), and expression of the ESX-3 operon is dependent on the transcriptional regulator IdeR, which controls iron metabolism (*25*) and is required for growth in the human pathogen *M. tuberculosis* (*26*). The ESX-3 operon is 67% identical between the non-pathogenic model organism *M. smegmatis* and the pathogenic *M. tuberculosis* over the 4354 amino acids of the ESX-3 operon. This high degree of conservation and important role in cell growth makes ESX-3 an important candidate for small molecule inhibition(*27*), as blockade of ESX-3 will inhibit virulence and kill a broad range of pathogenic mycobacteria.

To facilitate native purification of the translocon complex, a cleavable GFP tag was inserted into the chromosome of *M. smegmatis* MC^2^155 wild type and *ΔideR* (*28*) strains at the C-terminus of EccE_3_ via the ORBIT method (*29*) (Fig. 1A, Fig. S1). EccE_3_ is the final gene in the 11 gene long ESX-3 operon making insertion at this site less likely to disrupt regulation of the operon. Deletion of the iron acquisition regulator IdeR greatly increases chromosomal expression of ESX-3 (Fig. S2A). Four components of the complex EccB_3_, EccC_3_, EccD_3_ and EccE_3_ were affinity-purified as a large molecular weight species (Fig. 1B and 1C, Fig. S2B) using a GFP-nanobody and the GFP tag was proteolytically cleaved.

**Fig. 1.**
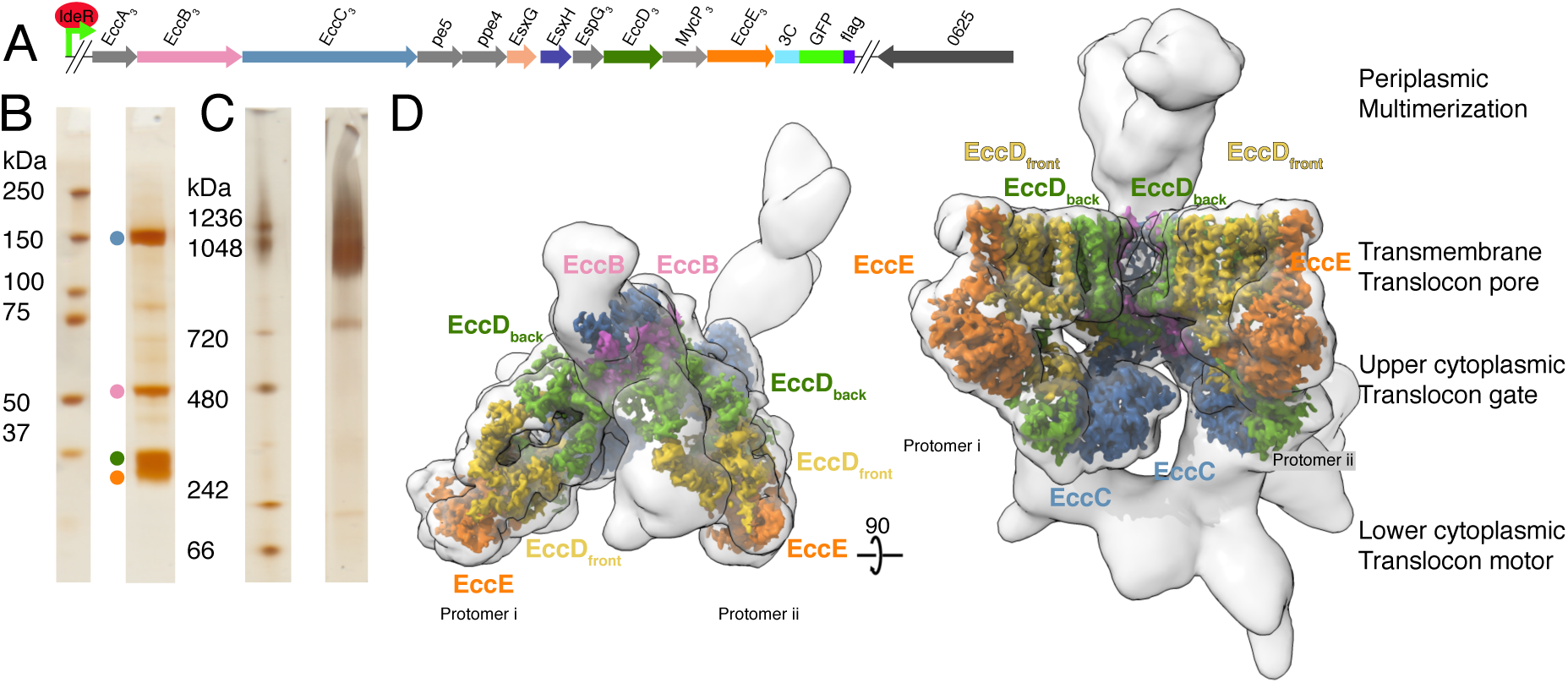
Overview of the ESX-3 tagging, purification, and structure. (A) The ESX-3 operon in *M. smegmatis* and the placement of the purification tag. Genomic deletion of *ideR* derepresses ESX-3 to boost expression for purification. (B) SDS-PAGE of purified ESX-3 shows four major bands corresponding to EccB_3_, EccC_3_, EccD_3_, and EccE_3_. (C) Blue native page of the purified ESX-3 complex shows a single large molecular weight band around 900 kDa. (D) Merged map of all focused refinement maps (grey transparency) of the ESX-3 translocon complex filtered to 10Å resolution. The transmembrane and upper cytoplasmic focused maps (3.7Å) segmented by subunit showing one copy per protomer of EccB_3_ (pink), EccC_3_ (blue), EccE_3_ (orange), EccD_3-front_ (yellow), EccD_3-back_ (green).

The ESX-3 translocon complex was imaged by cryo-EM and reconstructed revealing a dimeric structure which can be divided into four functionally important areas, the transmembrane translocon pore, the cytoplasmic translocon gate, the cytoplasmic motor domain, and the periplasmic multimerization domain (Fig. 1D, Fig. S3, Fig. S4, and Table S1). Particles of the size of a monomer or multimers larger than dimers were not observed in the electron microscopy data. The resolution of the ESX-3 translocon complex varies substantially in different parts of the electron microscopy map and this heterogeneity was partially resolved through data processing (see supplemental text, Fig. S5). The ESX-3 translocon complex is comprised of ten total proteins, two copies each of EccB_3_, EccC_3_, and EccE_3_ and four copies of EccD_3_. Two pseudo-symmetric protomers referred to as i and ii, combine to form the ESX-3 translocon complex. Each protomer contains one copy of EccB_3_, EccC_3_, and EccE_3_ and two conformationally distinct copies of EccD_3_, referred to as EccD_3-front_ and EccD_3-back_ (Fig. 2A). At 3.7Å resolution, the translocon pore and translocon gate were possible to build *de novo* atomic models for all observable amino acids except the two transmembrane helices of EccC_3_ (Fig. 2A, Table S2).

**Fig. 2.**
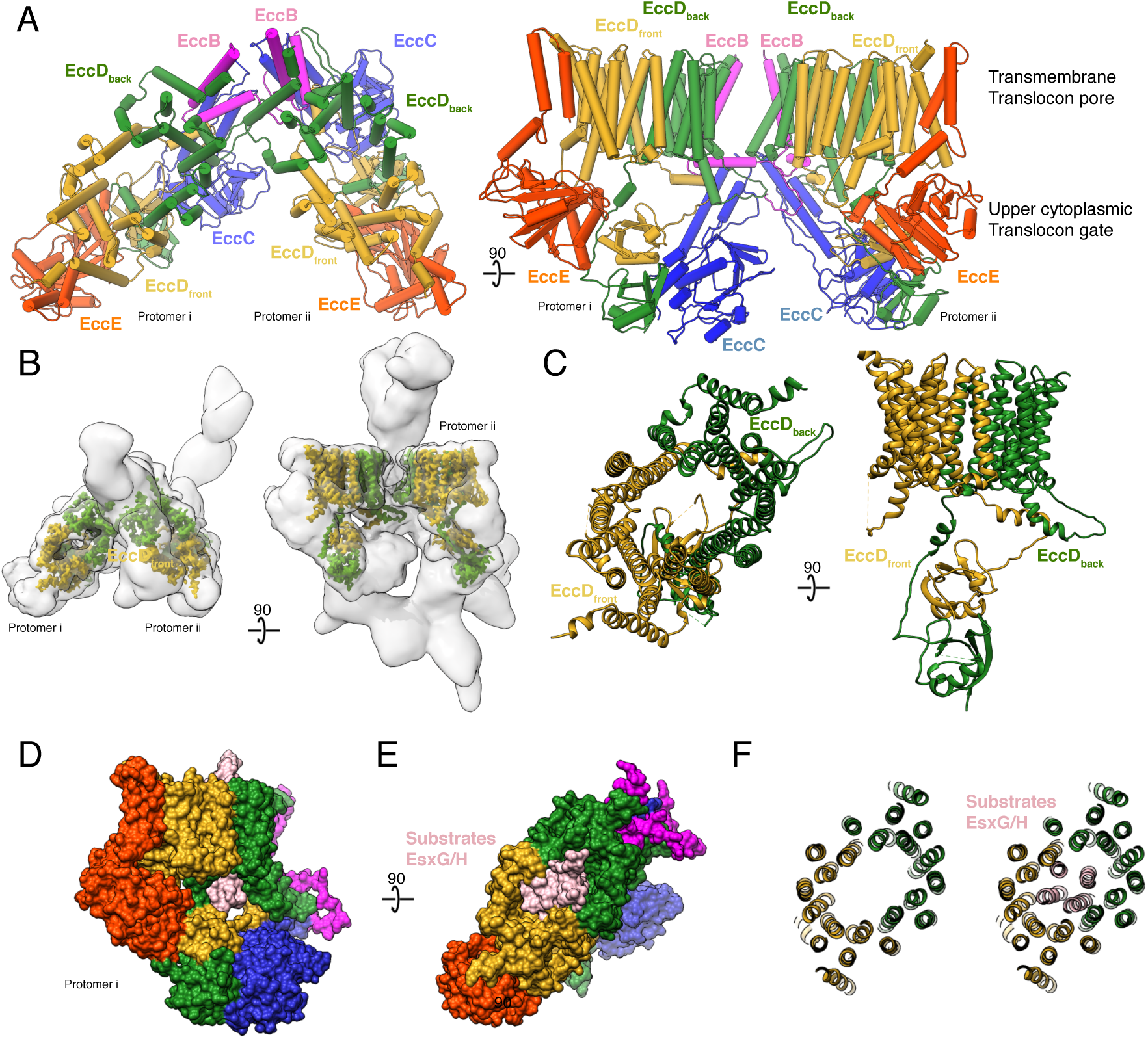
The translocon pore. (A) Atomic models of the translocon pore and gate. (B) EccD_3-front_ (yellow) and EccD_3-back_ (green) in the context of the overall ESX-3 translocon complex (grey transparency). (C) Atomic models of EccD_3-front_ and EccD_3-back_ (D) Space filling model of a single ESX-3 protomer with the substrate EsxG_3_-EsxH_3_ (PDB 3Q4H) (grey) fitted into the pore.

In each ESX-3 protomer, a distinct transmembrane pore is formed by the dimerization of the two copies of EccD_3_, EccD_3-front_ and EccD_3-back_ (Fig. 2B and 2C, Fig. S6A-C). The twenty-two EccD_3_ *α*-helices in each translocon pore form an elliptical, cylindrical void through the membrane with C2 symmetry and a cross-sectional diameter of ∼20 × 30 Å without significant regions of constriction (Fig. S6A). The inner surface of the translocon pores are composed primarily of hydrophobic residues (Fig. S6B) and in our maps, extended densities consistent with hydrophobic lipids or detergent molecules line the inner face of the pore (Fig. S6C). In contrast, the cytoplasmic face of the translocon pores are positively charged due to the presence of four basic residues (R134 and R239 from each EccD subunit) lining this region (Fig. S6B). Docking of the known structure of the dimeric EsxG_3_-EsxH_3_ (PDB 3Q4H) (*17*) substrate into the translocon pores reveals an opening sufficient to allow for the transit of the fully folded substrate monomers and dimers (Fig. 2D-F).

The translocon gate is formed by interactions between the cytoplasmic domains of EccD_3-_ front, EccE3, EccD3_-back_, and EccC_3_. In contrast to the translocon pore, the N-terminal, cytoplasmic domains of EccD_3_ are in strikingly different orientations, driven by alternative conformations of the residues joining the pore to the cytoplasmic domain (residues 100-127). This region of EccD_3_, adopts two distinct secondary structures resulting in the asymmetric placement of the cytoplasmic domains of EccD_3_ (Fig. 3A). In EccD_3-front_, residues 100-127 are bent, folding into an *α*-helix and forming extensive stabilizing contacts with EccB_3_ and EccC_3_ (Fig. S7A-G). In EccD_3-back_, residues 100-127 are extended and fold into a shorter *α*-helix that interacts with EccE_3_ and the cytoplasmic domain of EccD_3-front_ (Fig. S7H). This conformational flexibility suggests that residues 100-127 may act as a hinge that changes its conformation during the secretion cycle, facilitating the movement of substrates from the cytoplasm to the translocation pores or blocking transit through the pores.

**Fig. 3.**
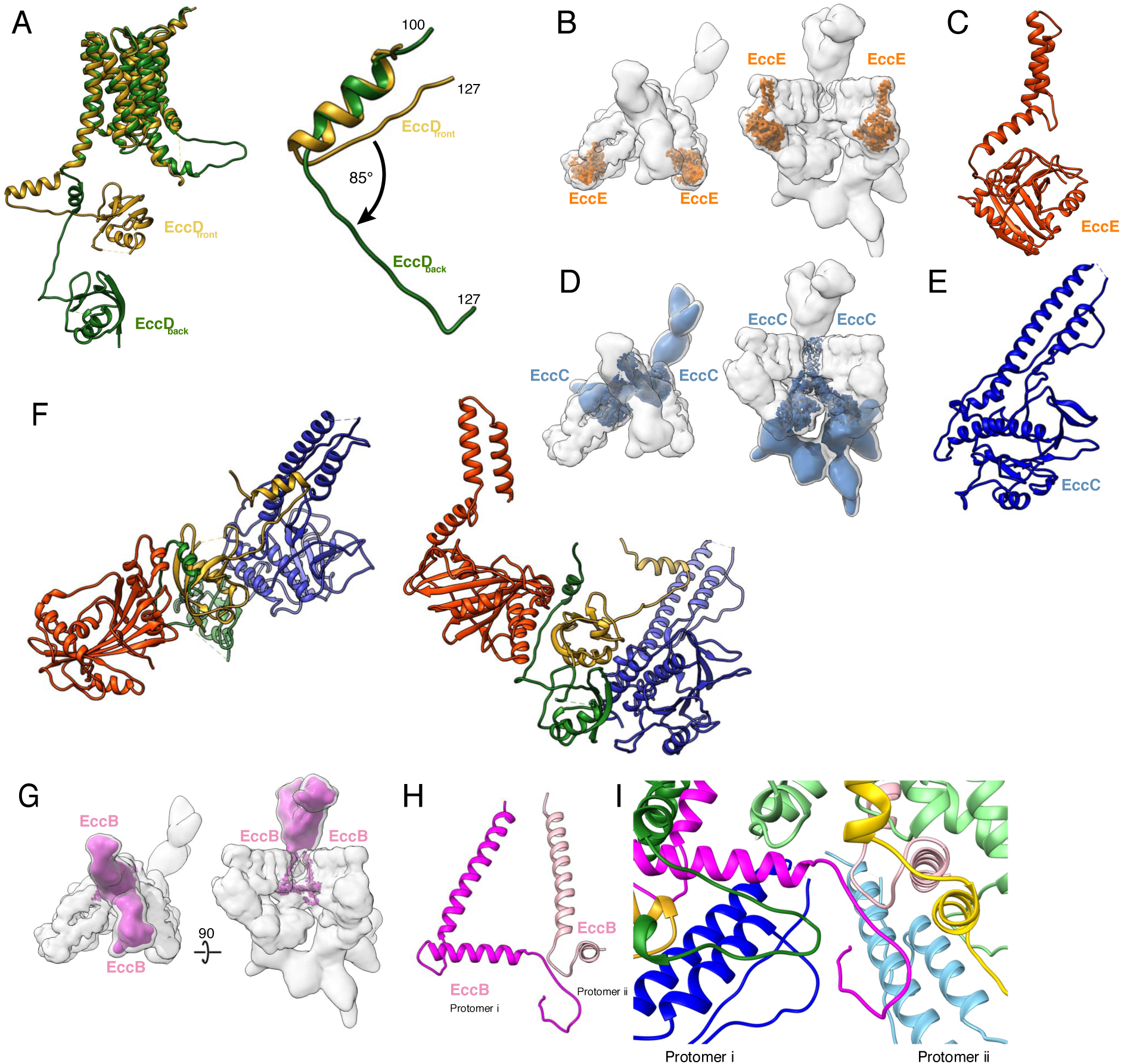
The translocon gate, motor, and multimerization domains. (A) EccD_3-front_ (yellow) and EccD_3-back_ (green) aligned based on the transmembrane regions shows two distinct conformations of the EccD_3_ cytoplasmic domains. Amino acids 100-127 of EccD_3_ adopt a bent (yellow) and an extended (green) conformation. (B) The placement of EccE_3_ in the overall ESX-3 translocon complex. (C) Atomic model of EccE_3_ (D) The overall placement of EccC_3_ in the ESX-3 translocon complex. EccC_3_ was reconstructed to variable resolutions. The upper cytoplasmic pseudoATPase domain (aa 1-37 and 94-403) in the translocation gate was reconstructed to 3.7Å. The lower cytoplasmic, ATPase 1, 2, and 3 domains in the translocon motor were reconstructed to ∼10Å resolution. (E) Atomic model of the EccC_3_ psuedoATPase domain. (F) Atomic model of the translocon gate comprised of EccE_3_ (orange), the cytoplasmic domain of EccD_3-front_ (yellow), the cytoplasmic domain of EccD_3-_ back (green), and the pseudoATPase domain of EccC_3_ (blue). (G) EccB_3_ (pink) in the context of the overall ESX-3 translocon complex (grey transparency). EccB_3_ has a single pass transmembrane domain which extends into a large periplasmic domain which was resolved at 5.5Å resolution. (H) Atomic models of the EccB_3_ cytoplasmic and transmembrane domains, amino acids 14-93 and 32-93. (I) The N-terminus of EccB_3_ forms extensive cross-protomer contacts with EccC_3_ (blue), EccD_3-front_ (yellow), and EccD_3-back_ (green).

The next component of the translocon gate is EccE_3_. EccE_3_ is positioned at the front of the ESX-3 translocation complex, interacting with helix 11 of EccD_3-front_ in the membrane and then extending into the cytoplasm (Fig. 3B and 3C, Fig. S8A-C). The cytoplasmic domain of EccE_3_ has weak structural homology to glycosyl transferase proteins, however, the nucleotide binding pocket is absent in EccE_3_ leaving it incapable of preforming this function (Fig. S8D and S8E, Table S4). Two helices in the cytoplasmic region of EccE_3_ between amino acids 133 and 163 form extensive stabilizing interactions with both subunits of EccD_3_ (Fig. S8F). These interactions hold the hinge region of EccD_3-back_ in the extended conformation and sterically hinder EccD_3-back_ from assuming the bent conformation seen in EccD_3-front_ (Fig. S8G).

The final component of the translocon gate is the domain of unknown fuction (DUF) of EccC_3_. EccC_3_ extends from the membrane into the upper and lower cytoplasmic regions (Fig. 3D). Amino acids 1-33 and 94-403 of EccC_3_ were built *de novo* into the higher resolution region of the electron microscopy map revealing the structure of the DUF domain (Fig. 3E, Fig. S9A-C). The *de novo* model of the DUF has the typical Rossman fold of a nucleotide hydrolysis domain, however, the Walker A motif contains substitutions and deletions that make ATP binding or hydrolysis by this domain unlikely (Fig. S9D and S9E). The protein binding helix remains intact and interacts with EccD_3back_ (Fig. S9F) suggesting the DUF functions as a pseudoATPase in EccC_3_ (*30*). The pseudoATPase domain is linked to the transmembrane domains by a long helical bundle making extensive contacts with both the hinge region of EccD_3front_ orienting the cytoplasmic domain of EccD_3front_ asymmetrically (Fig. S7G). Neither the pseudoATPase domain of EccC_3_ nor the cytoplasmic domain of EccE_3_ appear capable of binding and hydrolyzing nucleotides. Instead, these proteins appear to collaborate to buttress the cytoplasmic domains of EccD_3_ and form gates in the cytoplasm positioned to regulate the translocon pores (Fig. 3F).

The motor domain of the ESX-3 translocon complex contains the EccC_3_ ATPase 1, 2 and 3, which hang below the pseudoATPase domain, were resolved at low resolution ∼10Å, and are asymmetric between protomers i and ii (Fig. 3D, Fig. S9F and S9G). While both protomers have an identical protein composition, they differ in the conformations of their EccB_3_ and EccC_3_ components. When the translocation pore and gate are compared between protomers, only the transmembrane helices of EccC_3_ and the N-terminal tail of EccB_3_ differ (Fig. S10A and S10B). In contrast, the EccC_3_ ATPase 1, 2, and 3 domains do not super impose accurately across protomers even at low resolution (Fig. S10C-E). The heterogeneity of the motor domains suggests this region is dynamic throughout the translocation cycle.

The ESX-3 translocon complex dimer is stabilized by cross-protomer interactions formed by the two EccB_3_ proteins which begin in the cytoplasm with a flexible N-terminal tail leading into a linker helix, followed by a single pass transmembrane helix, and an extended periplasmic domain (Fig. 3G and 3H, Fig. S11). The N-terminal tail of EccB_3_ from protomer i forms extensive cross-protomer contacts with EccB_3_, EccC_3_, EccD_3front_ and EccD_3back_ from protomer ii (Fig. 3I, Table S3). The two EccB_3_ periplasmic domains share a large interaction interface across the protomers further stabilizing dimerization. Homology models of two EccB_3_ proteins can be docked into the periplasmic domain (Fig. S9C), however, this region is not resolved sufficiently to identify specific interactions. The majority of cross-protomer interactions involve EccB_3_ suggesting the periplasmic domain is essential for oligomerization.

The ESX-3 structure presented here is purified without the addition of substrates or nucleotide. It is therefore likely to be in a conformation representing the end of the translocation cycle, awaiting either the direct binding of substrates or the binding of nucleotide to reset a substrate-binding competent state. The EsxB substrate is known to be recruited either as a homo- or heterodimer in the cytoplasm by the ATPase 3 domain of EccC. Our structure is consistent with earlier reports that ESX substrates leave the translocation complex as folded dimers. The cytoplasmic domains of EccC, EccD, and EccE are idealy placed to regulate and guide substrates through the EccD pores (Fig. 4). The complexity of the ESX cytoplasmic assembly likely reflects the need for exquisite control of translocation in response to environmental changes and intracellular concentrations of substrates.

**Fig. 4.**
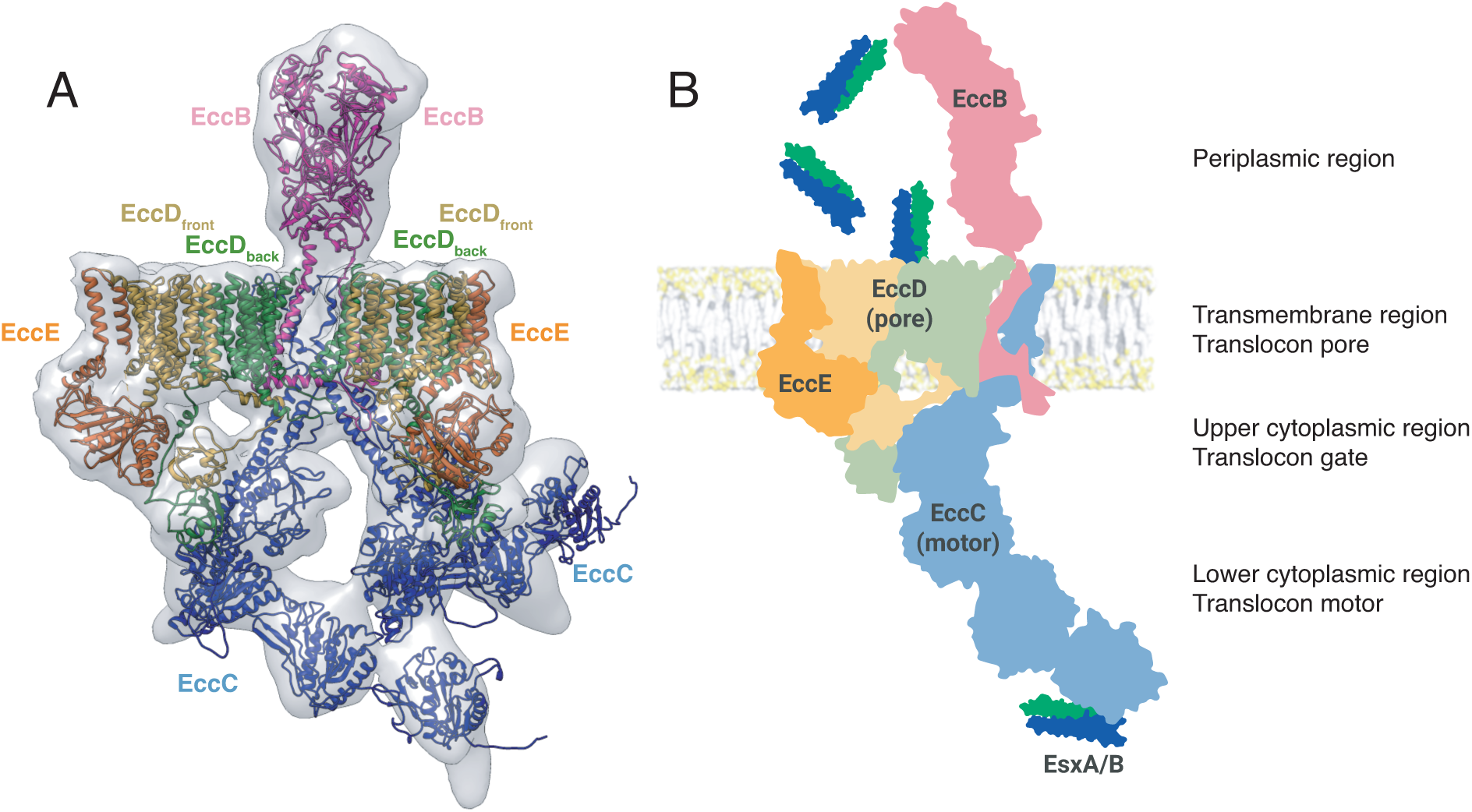
Hybrid model of the ESX-3 translocon complex. (A) A combined map of the full complex filtered to 10Å resolution (grey transparency) with models for each protein, EccB_3_ (pink), EccD_3-front_ (yellow), EccD_3-back_ (green), EccC_3_ (blue), and EccE_3_ (orange). (B) A model of the function of a single protomer of the ESX-3 complex. Substrates are selected by interaction with ATPase 3 of EccC and transported via the upper cytoplasmic gate to the translocon pore for secretion.

Each protomer contains all the components required to translocate substrates across the inner membrane: the transmembrane translocation pore (EccD), the cytoplasmic translocation gate (EccE, EccD, and EccC), and the cytoplasmic translocation motor domain (EccC). However, our structure of ESX-3 is clearly a dimer. The asymmetry of the EccC motor domains suggests one molecule of EccC may catalyze ATP hydrolysis in the second EccC molecule, thus making dimeric ESX-3 more active than single protomers alone. Previous reports have shown ESX-1 and ESX-5 form higher oligomeric states (*31*). Nothing in our structure prohibits additional ESX-3 protomers from joining together to form a higher order oligomeric state, and the unresolved flexible N-terminus of EccB is poised to interact with an additional protomer. Oligomerization may play an important role in spatial recruitment, regulation, and activation for targeted secretion of ESX substrates.

Our structure of a type VII secretion system provides a roadmap for the further exploration of a ubiquitous class of secretion systems found in most gram-positive bacteria and provides a scaffold for drug discovery targeting mycobacterial ESX translocon complexes.

## References and Notes

## Acknowledgments

The ΔIdeR strain was provided by Gabriella Marcela Rodriguez. Data was collected at the electron microscopy core facility at the University of California, San Francisco with the assistance of Alexander G. Myasnikov. We thank Robert Stroud and the members of his laboratory for stimulating discussions.

## Data and materials availability

Electron microscopy micrographs are deposited on EMPIAR. The electron microscopy maps are deposited to EMDB. The atomic models are deposited to the PDB.

## Supplementary Materials

### Materials and Methods

#### Strain construction

*Mycobacteria smegmatis* MC^2^155 wild type and ΔIdeR cells were chromosomally tagged using the ORBIT protocol (Fig S1). For wild type cells, the integrase expressing plasmid was pkm444. For ΔIdeR cells which already contained a kanamycin resistance marker, a zeomyocin resistance cassette was inserted into the pkm444 plasmid at the EcoIV restriction site. The tagging plasmid was pkm468 with a 3C insertion site added before the eGFP tag via primer extension PCR. The targeting oligo had the sequence: 5’ TGTGCGTTCCACTGGTTCCCCGGCAACCACCTGCTGCACGTGAGCCAGCCGGACTACCTAGGTTTGTACCGTACACCACTGAGACCGCGGTGGTTGACCAGACAAACCCGCCGGATGACCCGCTTCCTGCGCGGCTTCATGTTCGACTGAACCCTTCACCGAGGTCCG 3’. Bacterial cells with the integrase containing plasmid were grown in an overnight liquid culture and electroporated with a targeting oligo and tagging plasmid. The transformed *M. smegmatis* were plated on hygromycin containing 7H9 plates and incubated at 37° C for 3 days. Colonies were verified for insertion of the tagging plasmid into the chromosome by PCR.

#### Western Blotting

100mL of EccE_3_ tagged wild type and IdeR knock out cells were grown overnight to an OD of 1.0-1.2. Cells were pelleted and resuspended in 1 ml of buffer (50 mM Tris-HCl pH 8.0, 150 mM NaCl, 1% DDM) and sonicated for 30 seconds. Cell lysates were run on a 4-20% SDS-PAGE gel (Genescript) and transferred to blotting paper using a BioRad Trans-blot turbo system. The blot was washed with PBS and blocked in a 5% milk solution for 1 hour. The blot was incubated with mouse anti-GFP monoclonal antibody (Roche) overnight. After rinsing with PBS-T, the blot was incubated with anti-mouse IgG HRP-conjugated antibody (R&D Systems) for 2 hours. After activation (Amersham) the blot was imaged on a BioRad ChemiDoc. The blot was stripped with stripping buffer (ThermoFisher Scientific) as per the manufactures instructions, and incubated overnight with rabbit anti-GroEL monoclonal antibody (Sigma-Aldrich). The blot was incubated with goat anti-rabbit IgG antibody HRP (GenScript) for 2 hours, activated (Amersham), and imaged on a BioRad ChemiDoc.

#### Protein purification

*M. smegmatis* was grown in 7H9 supplemented with 0.05% Tween 80 and 20 µg/ml kanamycin. After harvest, cells were washed three times with PBS and frozen in liquid nitrogen before lysis with a cryogenic grinder (SPEX SamplePrep). 24.9g of powdered cell material was resuspended in 56.3 ml 50 mM Tris-HCl pH 8.0, 150 mM NaCl, 1% DDM with protease inhibitor cocktail (SigmaFast) and 175 units Benzonase endonuclease. The suspension was stirred for 120 min at 4°C. After centrifugation for 30 min at 98,000 g (ThermoLynx), the supernatant was incubated with 1.4 ml anti-GFP-nanobody resin for 90min at 4°C. The resin was transferred to a column and washed sequentially with 28ml of wash buffer (50 mM Tris-HCl pH 8.0, 150 mM NaCl and 0.1% GDN), 14ml of high salt wash buffer (50 mM Tris-HCl pH 8.0, 400 mM NaCl, and 0.1% GDN), and 14ml of wash buffer (50 mM Tris-HCl pH 8.0, 150 mM NaCl, and .1% GDN). To cleave off the purification tag, the resin was incubated o/n at 4°C with 70 units Pierce HRV 3C protease (ThermoFisher Scientific) in 2.8 ml wash buffer supplemented with 0.2 mM DTT. This resin was sedimented by gentle centrifugation (300 × g for 3 min), the supernatant collected, and the resin was subsequently washed with 1.4 ml wash buffer. The supernatant and first wash fraction were combined and concentrated using an Amicon Ultra-4 centrifugal filter unit (100 kDa molecular weight cut-off). The sample was centrifuged for 6 min at 16,000 g before injection on a Superdex 200 (10/300) column equilibrated in 50 mM Tris-HCl pH 8.0, 150 mM NaCl and 0.021% GDN. Peak fractions were concentrated using a 0.5 ml centrifugal filter unit (Amicon, 100 kDa cut-off) to 5.52 mg/ml by Nanodrop reading.

#### Blue-Native Polyacrylamide Gel Electrophoresis (BN-PAGE)

BN-PAGE was carried out using the Invitrogen NativePAGE Bis-Tris Gel system. Samples were prepared in a total volume of 10 µl using 0.25µl 5% G-250 sample additive. Electrophoresis was performed using light blue cathode buffer at a constant voltage of ∼ 105 V for 3.5 h at 4°C. The gel was fixed and stained using the Pierce silver stain kit.

#### Cryo electron microscopy – data acquisition

Samples were frozen for cryo electron microscopy. Quantifoil R1.2/1.3, 400 mesh, cooper grids were glow discharged using a Solarus plasma cleaner (Gatan) with an H_2_/O_2_ mixture for 30 seconds. 2µl of sample were applied per grid and the grids were plunged into liquid ethane using a FEI Vitrobot Mark IV.

Initially, samples were screened, and test data sets were collected on a FEI Talos Arctica 200kV microscope equipped with a Gatan K2 Summit detector. For the initial screen of freezing conditions, 834 movies were collected at a magnification of 36,000 with a pixel size of 1.14, and a defocus range of -1.5 to -2.5 µm, an exposure time of 9 seconds, and a dose rate of 7 electrons/Å^2^/second. (Table S1). Data collection for the final structure presented in the main text was collected on a FEI Titan Krios at 300kV with a Gatan K2 Summit detector. Two imaging sessions were used. In the first imaging session, 2705 movies were collected at a magnification of 29,000 with a pixel size of 0.82, and a defocus of -0.4 to -1.2 µm, an exposure time of 10 seconds to collect 100 total frames, and a dose of 8 electrons/Å^2^/second. (Table S1). In the second imaging session, data was collected on the same microscope with the same detector, 4632 movies were collected at a magnification of 29,000 with a pixel size of 0.82, and a defocus range of -0.6 to -1.4 µm, an exposure time of 10 seconds to collect 80 total frames, and a dose of 6.7 electrons/Å^2^/second.

#### Cryo electron microscopy – data processing

For all data, movies were motion corrected using MotionCor2(*1*) and CTF correction was performed using CTFfind4(*2*). For the Arctica dataset, particles were picked using a gaussian blob in either RELION(*3*) or cisTEM(*4*) and initial 2D classification was performed to remove obvious junk particles. Initially, a shotgun approach was taken to generate several initial models using RELION, cisTEM, and cryosparc(*5*). Once an initial model which contained realistic low-resolution features was generated, a user defined descent gradient was performed to improve the model with the goal of achieving accurate secondary structure features. First, all particles selected during 2D classification were refined in 3D against the randomly generated initial model. Second, a round of 3D classification with 4 classes and default RELION settings was performed and the best class selected. Third, the best class was refined as a single class in 3D classification with increasing Tau2_Fudge and decreasing search angle size. The resulting EM density map had clear transmembrane helix densities and was used as the model for a new 3D reconstruction. This reconstruction was used to back project models for reference-based particle picking in RELION. Two rounds of 2D classification were performed and the best classes selected. One round of 3D classification was performed using the Tau2_Fudge value optimized during the previous run through (T=12) and the best class selected. A final 3D reconstruction of the Arctica data set yielded a map of about 4.7Å resolution (Fig. S9).

After motion correction and CTF determination, the final Titan Krios dataset was processed entirely using RELION. Particles were picked using a gaussian blob, and extracted as 4x binned particles. Two rounds of initial 2D classification were performed with T=3 on the binned particles and obvious junk particles were removed. The final reconstruction from the Arctica dataset was used as the initial model for a 3D reconstruction of the binned particles. 3D classification with 4 classes and the previously optimized Tau2_Fudge value, T=12, was performed on the binned particles. The two best class were selected and re-extracted without binning. A 3D reconstruction was performed. A mask was created for the high-resolution region of the reconstruction and 3D classification without image alignment was performed focused on this region. The best class was selected and the subsequent 4.0Å reconstruction is the consensus structure for the entire complex (Fig. S3). Focused classification of each protomer, the periplasmic EccB region, and the ATPase 1,2, and 3 domains of EccC were performed. To perform focused classification, the center of mass of the region of interest was determined using chimera(*6*). Particles were recentered on this area and reextracted. Masks for the region of interest were generated and 3D classification without image alignment was performed. The best class was selected and used for a focused 3D reconstruction without image alignment of the region of interest. A reconstruction was generated and density outside of the region of interest was subtracted. A final reconstruction of the masked and density subtracted particles was then performed. This procedure improved the resolution of the protomers to 3.75Å (left) and 3.83Å (right), 5.6Å resolution for the EccB periplasmic domain, and ∼10Å resolution for the EccC lower cytoplasmic region.

To generate the symmetry expanded protomers based on non-point group symmetry (also known as non-crystallographic symmetry or NCS), a transformation matrix between the two protomers was calculated using chimera. Particles were then transformed and aligned using the subparticles.py and star.py utilities in pyem[cite] resulting in a particle stack with twice as many particles as the input file, each focused on protomer i or protomer ii. Density subtraction was performed to remove density outside of the symmetry expanded protomer, and focused classification and refinement were performed as described above. This procedure improved the resolution of the symmetry expanded protomer to 3.69Å resolution.

#### Atomic model building

The cytoplasmic domain from the crystal structure of EccD_1_ (PDB 4KV2) was docked into the cytoplasmic domains of the two EccD_3_ molecules and the sequence was mutated. The remaining transmembrane domains of EccD_3_ and the residues 14-93 of EccB_3_ were built *de novo* in Coot(*7*) using baton building. The alpha helices of EccE_3_ and EccC_3_ were initially modeled using the RaptorX(*8*) homology server. The loops and strands of EccE_3_ and EccC_3_ were built in Coot using baton building. All models were subsequently refined individually, as a symmetry expanded protomer, left and right protomers, and as the full model using phenix real space refine(*9*), Coot, and the MDFF(*10*) server, namdinator(*11*) (Table S2).

#### Low resolution modeling

The left and right protomer map, periplasmic focused refined map, and lower cytoplasmic focused refined map were all docked into the consensus map and added together using chimera. The combined map was filtered to 10Å resolution to match the lowest resolution component. Homology models for amino acids 94-516 of EccB_3_, the transmembrane helixes of EccC_3_, and 404-1268 of EccC_3_ were generated using RaptorX. These models were fit into the combined map density using the fit map to model utility in Chimera. The full model was refined using phenix.real_space_refine.

#### Model interpretation and display

The pore diameter and properties were analyzed using MOLE(*12*). Buried surface area between subunits was calculated by PISA(*13*). Atomic models for individual proteins were compared against the PDB using the DALI server(*14*). Chimera and chimerax(*15*) were used to display maps and models for figure creation.

### Supplementary Text

Initially, the entire ESX-3 translocon complex was reconstructed to 4.0Å resolution (Fig. S5A). Using symmetry expansion, and focused classification and refinement techniques, the resolutions of targeted regions of the ESX-3 complex were improved to 3.7Å for the transmembrane region and upper cytoplasmic region (Fig. S5B-D), 5.5Å resolution for the periplasmic region (Fig. S5E), and 10Å resolution for the lower cytoplasmic region (Fig. S5F). The highest resolution maps for each region were combined and filtered to the threshold of the lowest resolution map to form an overall 10Å consensus map for the entire translocon complex

The front EccD molecule forms the following interactions: Arg11 of the front EccD forms an ionic interaction with Glu161 of EccE. Asp14 of the front EccD forms an ionic interaction with Arg99 of EccE. Arg23 of the front EccD forms an ionic interaction with Asp111 of the back EccD. Glu26 of the front EccD forms an ionic interaction with Asp157 of EccE. Glu32 of the front EccD forms backbone contacts with Gly18 and Asp19 of the back EccD. Arg36 of the front EccD interacts with Glu389 of EccC. Pro40 of the front EccD interacts with His392 of EccC. Glu37 of the front EccD interacts with Arg395 of EccC. Arg66 of the front EccD interacts with Glu134 and Glu389 of EccC. Arg107 of the front EccD interacts with Arg116 of EccC. Glu 110 of the front EccD interacts with Arg112 of EccC. Asp111 and Asp114 of the front EccD interact with Arg47 of EccB. Phe119 of the front EccD interacts with Pro321 of the back EccD. Ser120 of the front EccD interacts ionically with Gln328 of the back EccD and His55 of EccB. Glu121 of front EccD interacts ionically with His55 of EccB. Arg123 of the front EccD interacts with Arg239 and Arg325 of the back EccD. Arg124 of the front EccD interacts with Asp378 of the back EccD and His55 of EccB. Arg125, His131, and Arg134 of the front EccD are coordinating a non-protein density together. Gln126 of the front EccD is interacting with Asp378 of the back EccD. Trp127 of the front EccD has a cation-π interaction with Lys383 of the back EccD. Pro129 of the front EccD has an interaction with Pro431 of the back EccD. The residues in transmembrane helix 1 of the front EccD have several non-specific hydrophobic interactions with transmembrane helices 9 and 10 of the back EccD. These last three interactions between the EccD front and back protein’s Trp127 and Lys383, Pro129 and Pro431, and transmembrane helix 1 and transmembrane helices 9 and 10 are mirrored on the other side of the pore. Transmembrane helix 11 of the front EccD forms non-specific hydrophobic interactions with transmembrane helix 1 of EccE.

The back EccD forms the following interactions: Arg11 of the back EccD protrudes into a negatively charged pocket on EccC formed by Asp268, Ser398, and Asp401. Gly18 and Asp19 of the back EccD have backbone interactions with Glu32 of the front EccD. Glu26 of the back EccD forms an ionic interaction with Arg395 of EccC. Asp92 of the back EccD interacts with Arg399 of EccC. Ala105 of the back EccD forms a backbone contact with Glu161 of EccE. Glu110 of the back EccD forms an ionic contact with Arg138 and Arg154 of EccE. Between Ile108 and Ile112 of the back EccD is a beta strand which complements the beta sheet of the cytoplasmic domain of the front EccD. Asp111 of the back EccD forms an ionic interaction with Arg23 from the front EccD.

Of the 80 amino acids modeled in EccB, almost all are involved in important inter or intra molecular interactions. Arg16 of EccB forms ionic contacts with Asp178 and Glu179 of EccC from the same protomer and could potentially from an ionic contact with Tyr107 of EccC from the opposite protomer. Thr17 of EccB forms an ionic contact with Arg106 of EccC from the opposite protomer. Asn20 of EccB forms an ionic contact with Asp113 of EccC from the opposite protomer. Glu21 of EccB forms an ionic contact with Asn114 of EccC from the opposite protomer. Asn22 of EccB forms an ionic contact with Asp113 of EccC from the opposite protomer. Asp24 of EccB forms an ionic contact with Arg116 of EccC from the opposite protomer. The side chain position of Arg29 of EccB is ambiguous and it could form an ionic contact with Asp306 of the back EccD molecule from the same protomer or Asp378 of the back EccD molecule from the opposite protomer. The side chain position of Arg30 of EccB is ambiguous and it could form ionic contact with Asp378 or Asp437 of the back EccD molecule from the opposite protomer. Phe32 of EccB forms a cation-π interaction with Arg375 of the back EccD from the opposite protomer. Arg35 of EccB forms an interaction with Arg445 of the back EccD from the opposite protomer. His36 of EccB might form an ionic interaction with Asp181 of EccC from the same protomer. Gln37 of EccB forms an ionic interaction with Asp306 from the back EccD from the same protomer. Ser39 of EccB forms an ionic interaction with Arg101 of EccC from the same protomer. Trp41 of EccB forms CH/π interactions with Pro442 and Pro229 from the back EccD from the same protomer. Arg42, Arg46, Arg65, and Arg69 of EccB all interact with each other to stabilize the loop between the linker and transmembrane helix of EccB. All four of these arginine residues may also interact with Asp98 of EccC from the same protomer. This interaction may also be involved in coordinating lipids or locally melting the membrane to allow for flexibility. Phe43 of EccB forms a cation-π interaction with Arg101 of EccC from the same protomer. Arg47 of EccB forms ionic contacts with Asp111 and Asp114 from the front EccD from the same protomer. His55 of EccB sits in a pocket of charged amino acids and participates in a complex interaction network with Glu121 and Arg124 from the front EccD and Asp378 from the back EccD all from the same protomer. Arg58 of EccB interacts with Arg106 of EccC from the same protomer. The transmembrane helix of EccB, Pro63 to Phe91, forms several non-specific hydrophobic interactions with the back EccD transmembrane helix 11 in the same protomer. Overall, EccB forms a substantial number of protein-protein interactions and all substantial cross-protomer interactions involve EccB making it essential for multimerization of ESX-3 protomers.

**Fig. S1.**
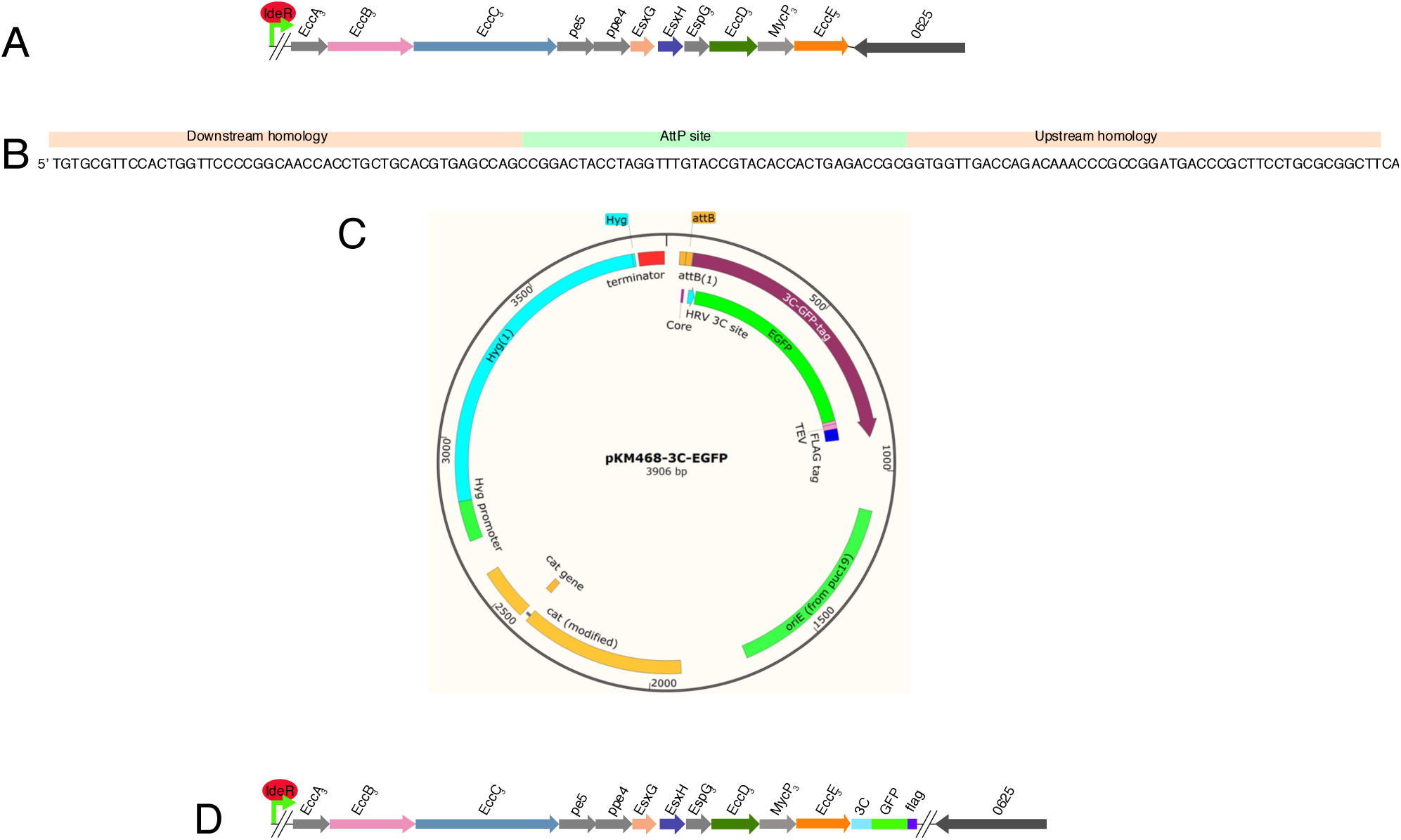
ORBIT tagging of the chromosomal copy of EccE_3_. (A) Cartoon of the ESX-3 operon present in the *M. smegmatis* MC^2^155 chromosome. (B) The targeting oligo used to insert the attp site into the chromosome. (C) The tagging plasmid recombined into the chromosome. (D) Cartoon of the resulting change to the chromosome after the ORBIT protocol was performed.

**Fig. S2.**
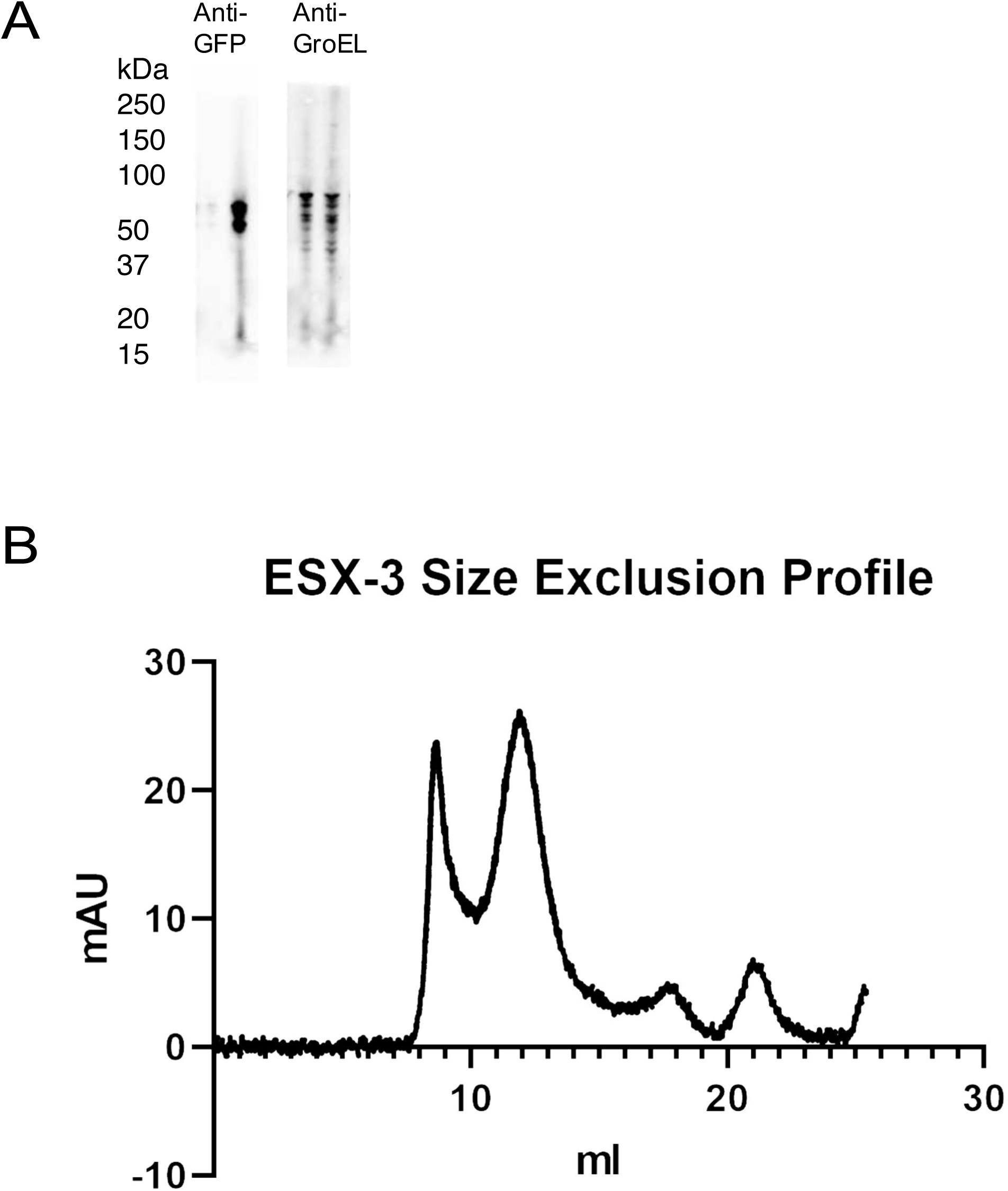
ESX-3 expression and purification. (A) Western blot of solubilized cell material with EccE3 tagged with GFP. Lane 1, wild type background. Lane 2, ΔIdeR background. (B) Size exclusion profile for the ESX-3 purification.

**Fig. S3.**
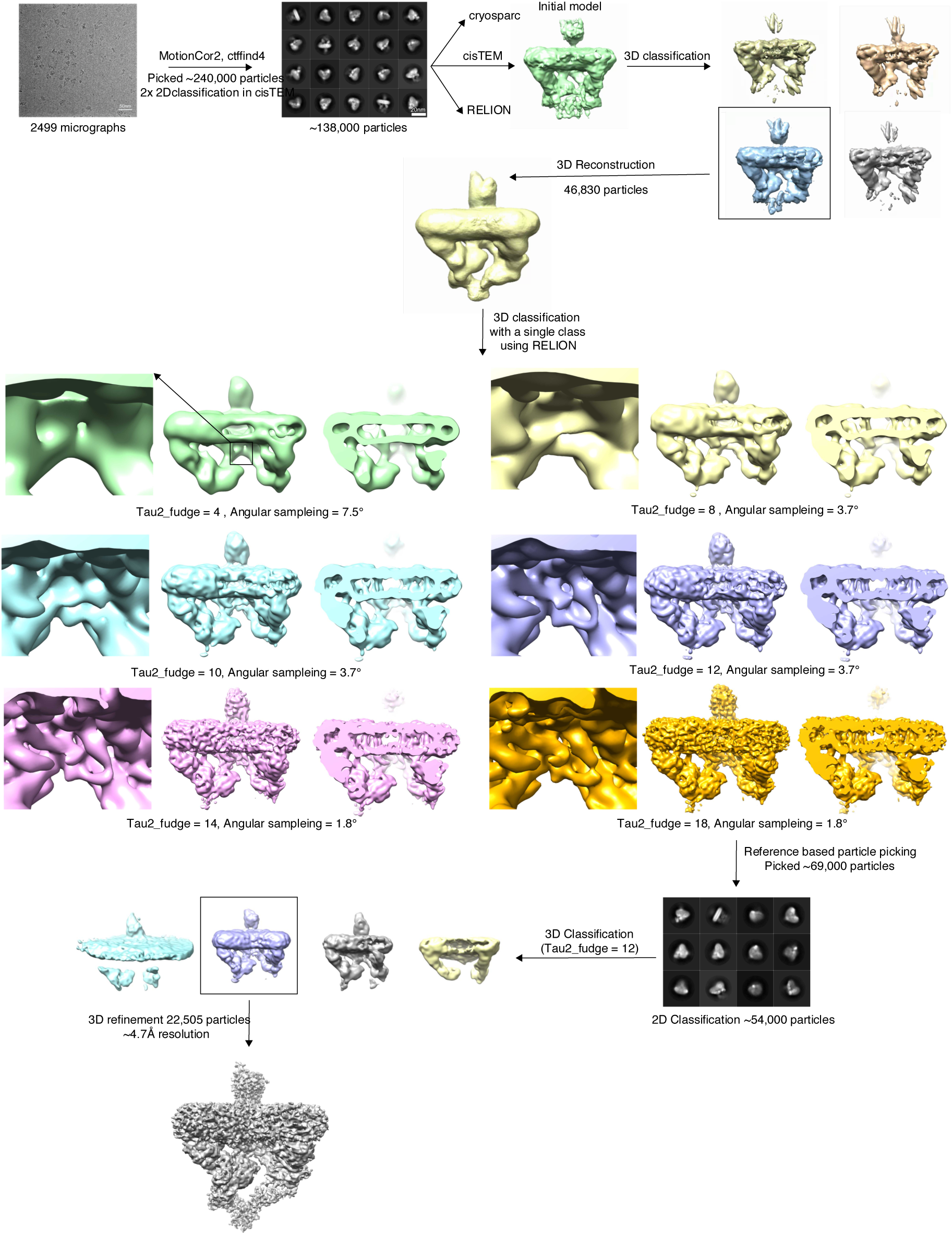
Initial data collection and initial model generation. An initial data set was collected on a Talos arctica microscope. The final refinement from this data processing was used as the starting model for future data processing

**Fig. S4.**
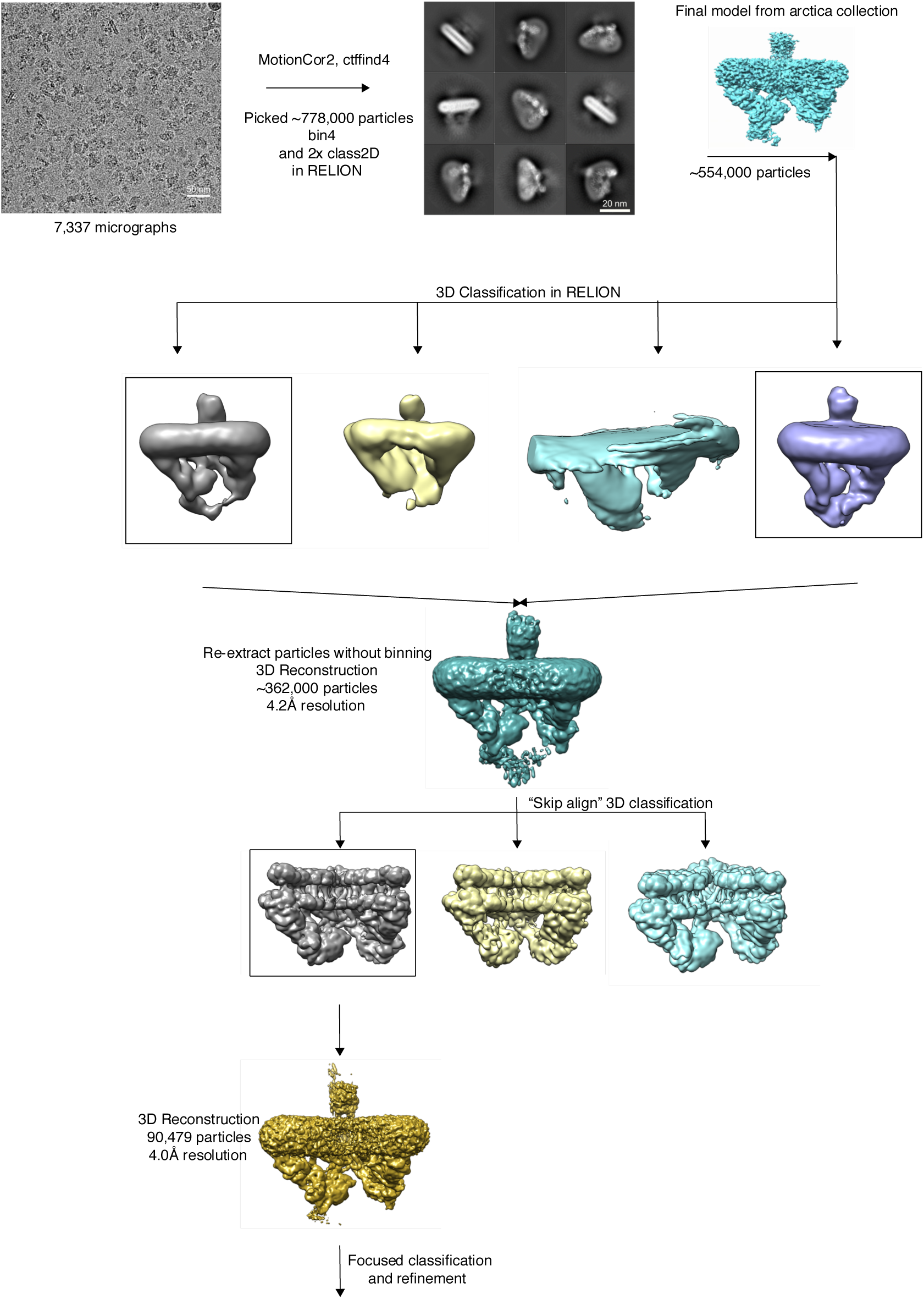
Data processing workflow for final data collection. Data was collected on a Titan Krios. Particles from the final reconstruction were subsequently used for focused classification and refinement.

**Fig. S5.**
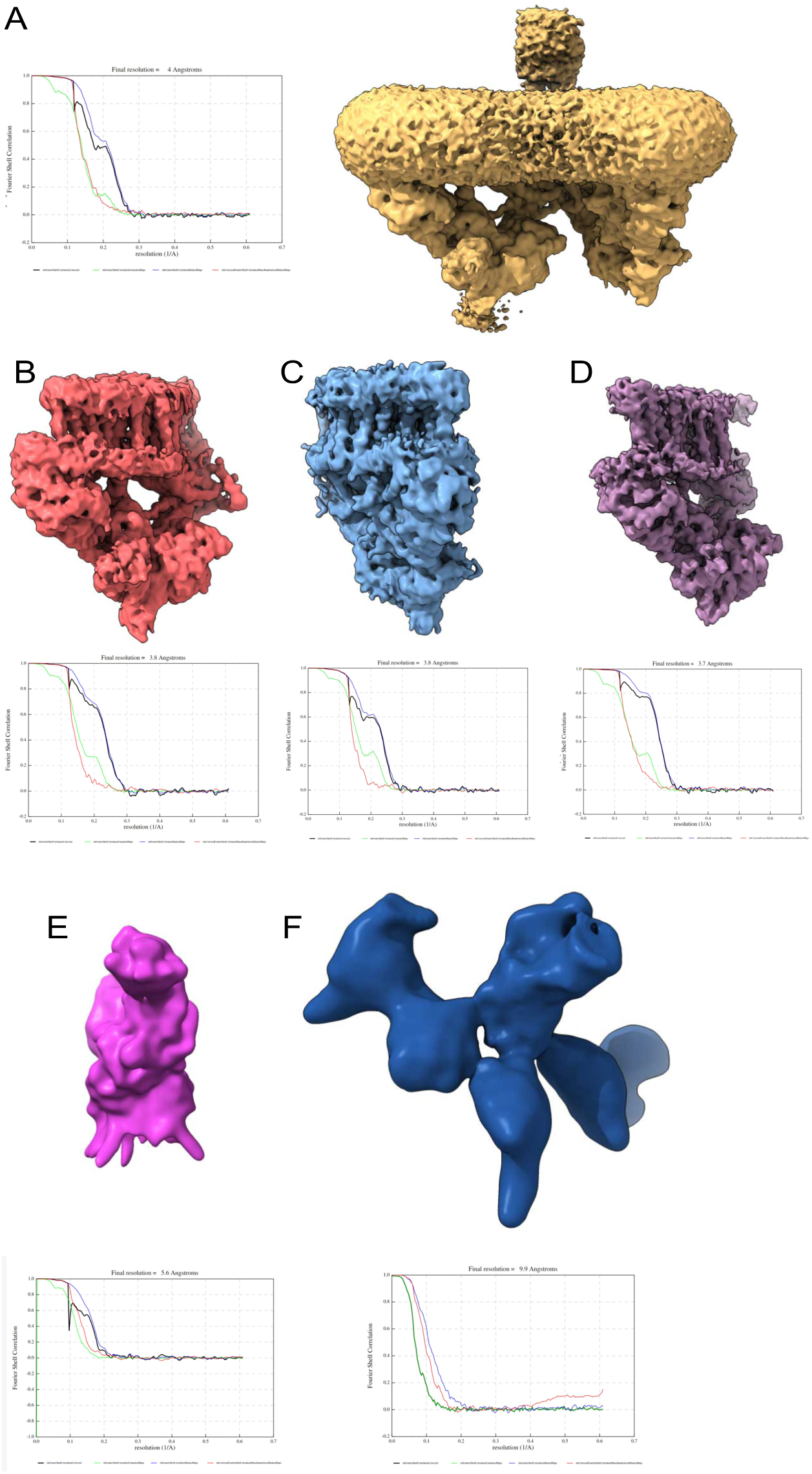
Consensus and focused refinements. (A) Consensus refinement map and FSC curve for the ESX-3 complex. Focused refinement maps and FSC curves for (B) the left protomer of the ESX-3 complex. (C) the right protomer of the ESX-3 complex (D) a symmetry expanded protomer (E) the periplasmic domain (F) the lower ATPase domains of EccC. All maps are unsharpened.

**Fig. S6.**
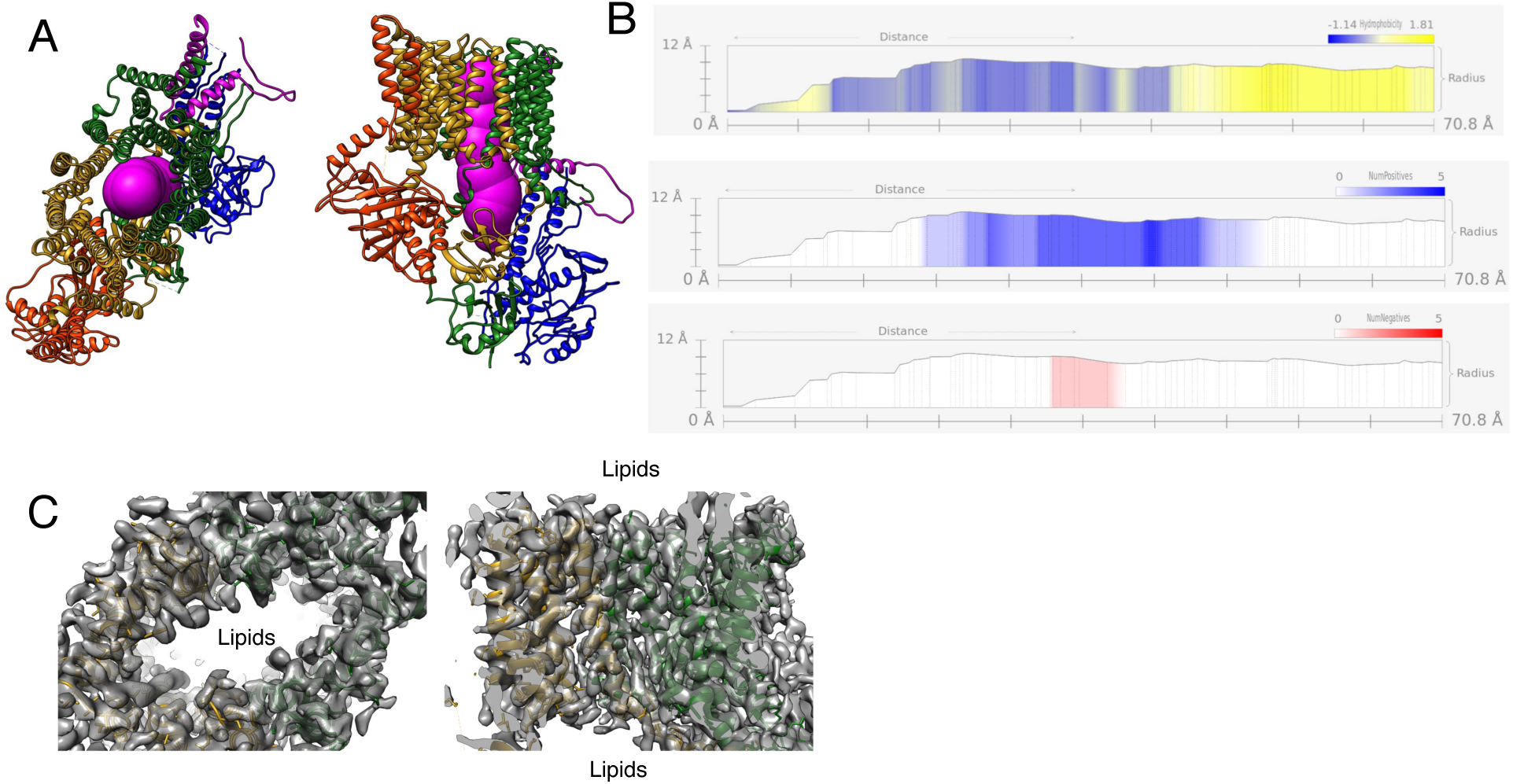
The translocon pore. (A) The path of the pore as mapped by the program MOLE shown in purple spheres through one protomer. (B) The path of the pore shows no constriction and positively charged at the cytoplasmic face. (C) Lipid and detergent line the inside of the pore.

**Fig. S7.**
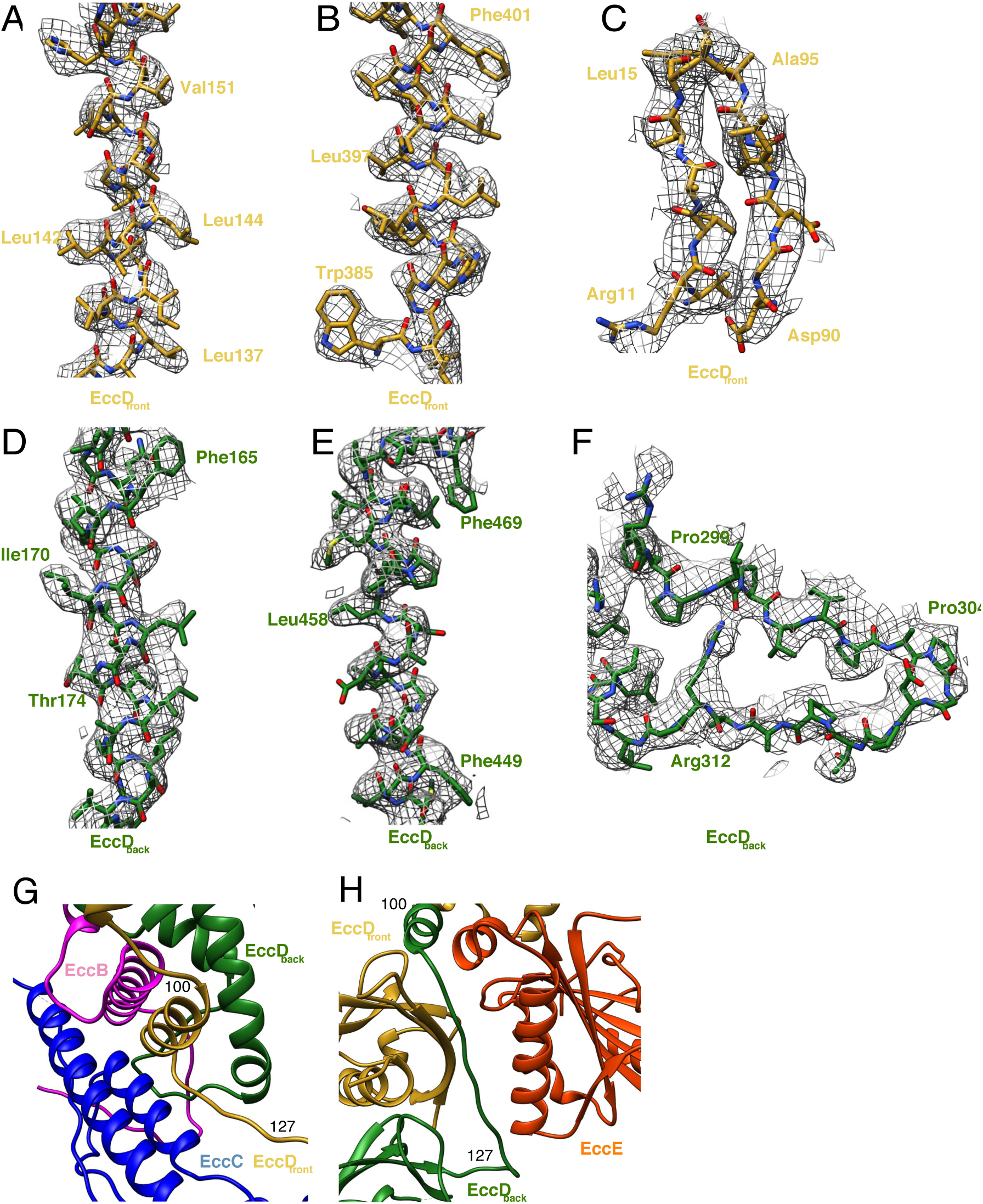
EccD_3_ map and model. Map to model fits for EccD_front_ for (A) transmembrane α-helix 1, amino acids 136-153, (B) transmembrane α-helix 1, amino acids 385-402, (C) soluble domain β-strands amino acids 10-15 and 90-96. Map to model fits for EccD_back_ for (D) transmembrane α-helix 1, amino acids 163-181 (E) transmembrane α-helix 11, amino acids 446-469 (F) soluble loop, amino acids 297-315. (G) Amino acids 100-127 from EccD_3-front_ (yellow) interact with EccB_3_ (pink) and EccC_3_ (blue). (H) Amino acids 100-127 of EccD_3-back_ (green), in a distinct conformation, interact with EccE_3_ (orange) and EccD_3-front_ (yellow)

**Fig. S8.**
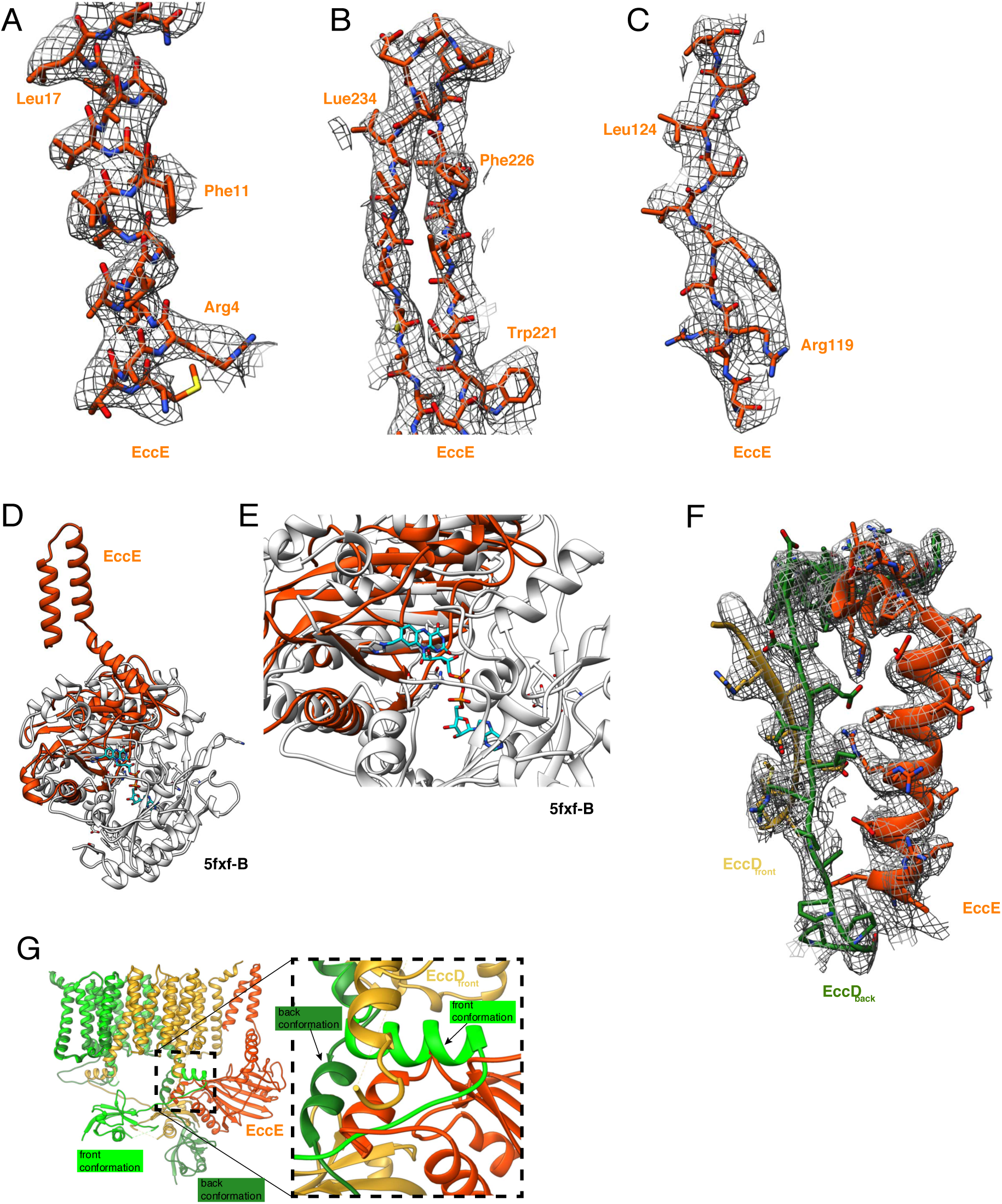
EccE_3_ map and model. (A) Map and model comparison for transmembrane helix 1, amino acids 1 to 18. (B) Beta strand separation, amino acids amino acids 221 to 240. (C) a single beta strand, amino acids 117 to 126. (D) Comparison between the structure of EccE_3_ and the top DALI hit 5fxf-B. (E) Close up on the FADH (cyan) binding pocket of 5fxf-B which is not present in EccE_3_. (F) Residues 133-163 of EccE_3_ interact substantially with EccD_3-front_ and EccD_3-back_, holding the hinge of EccD_3back_ in the extended confromation (G) EccE_3_ (orange) stabilizes the extended hinge conformation of EccD_3-_ back (green) and sterically hinders it from adopting the bent conformation (neon green).

**Fig. S9.**
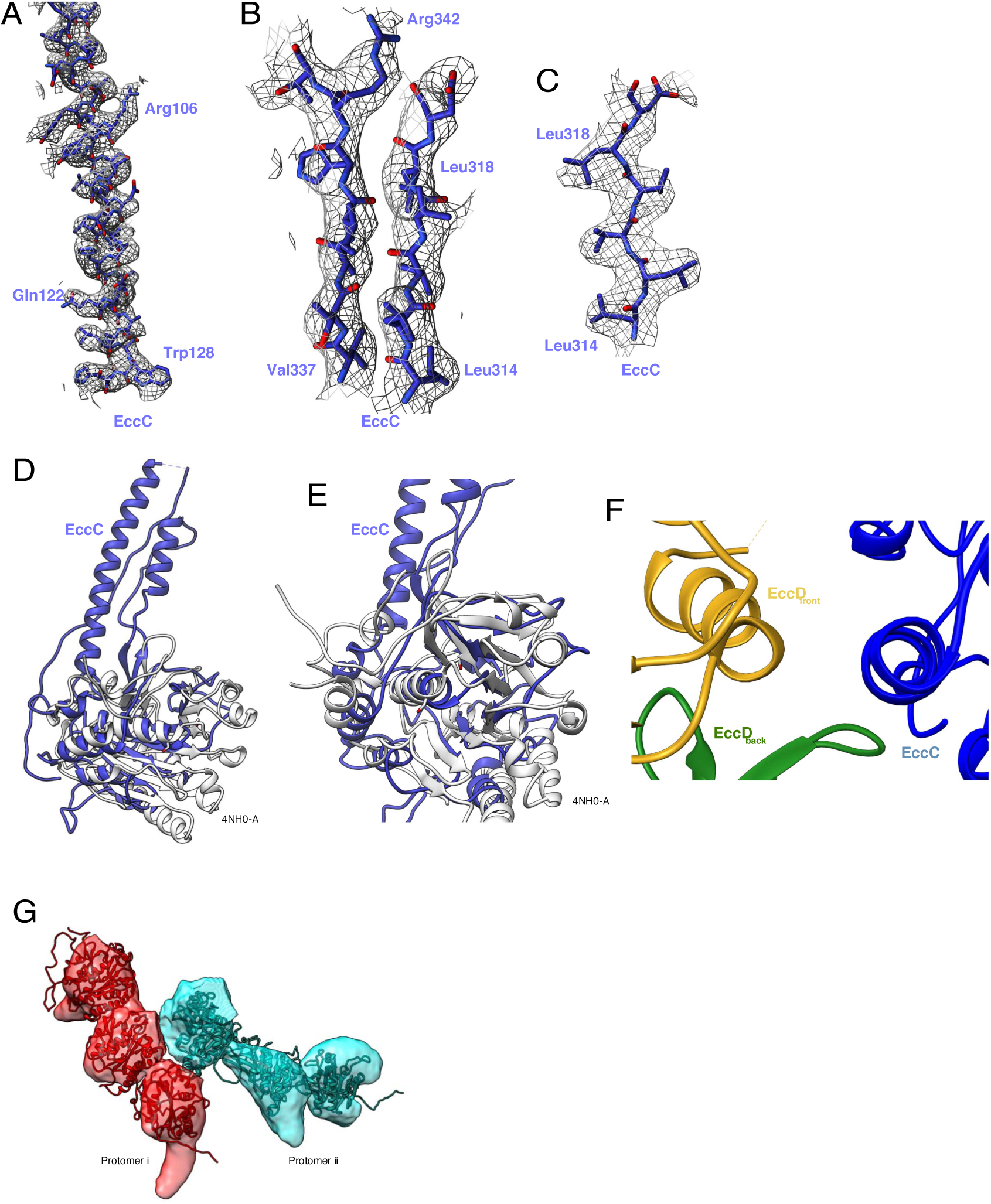
EccC_3_ map and model. (A) Map and model of the soluble helix of the EccC pseudo ATPase domain, amino acids amino acids 97 to 130. (B) Two beta strands in the EccC pseudo ATPase domain, amino acids 314 to 319 and 337 to 343. (C) Beta strand in the EccC pseudo ATPase domain, amino acids 314 to 319. (D) Overlay of the model of the EccC pseudo ATPase domain with the Top Dali hit, 4NH0-A (white). (E) Close up of the ATP binding pocket of 4NH0-A. While the fold remains intact in the pseudo ATPase domain, the amino acids required for ATP binding and hydrolysis are mutated. (F) The protein binding domain of EccC_3_ interacts with the cytoplasmic domain of EccD_3-front_ (yellow). (G) ATPase 1,2, and 3 domains of EccC_3_ with homology models docked in.

**Fig. S10.**
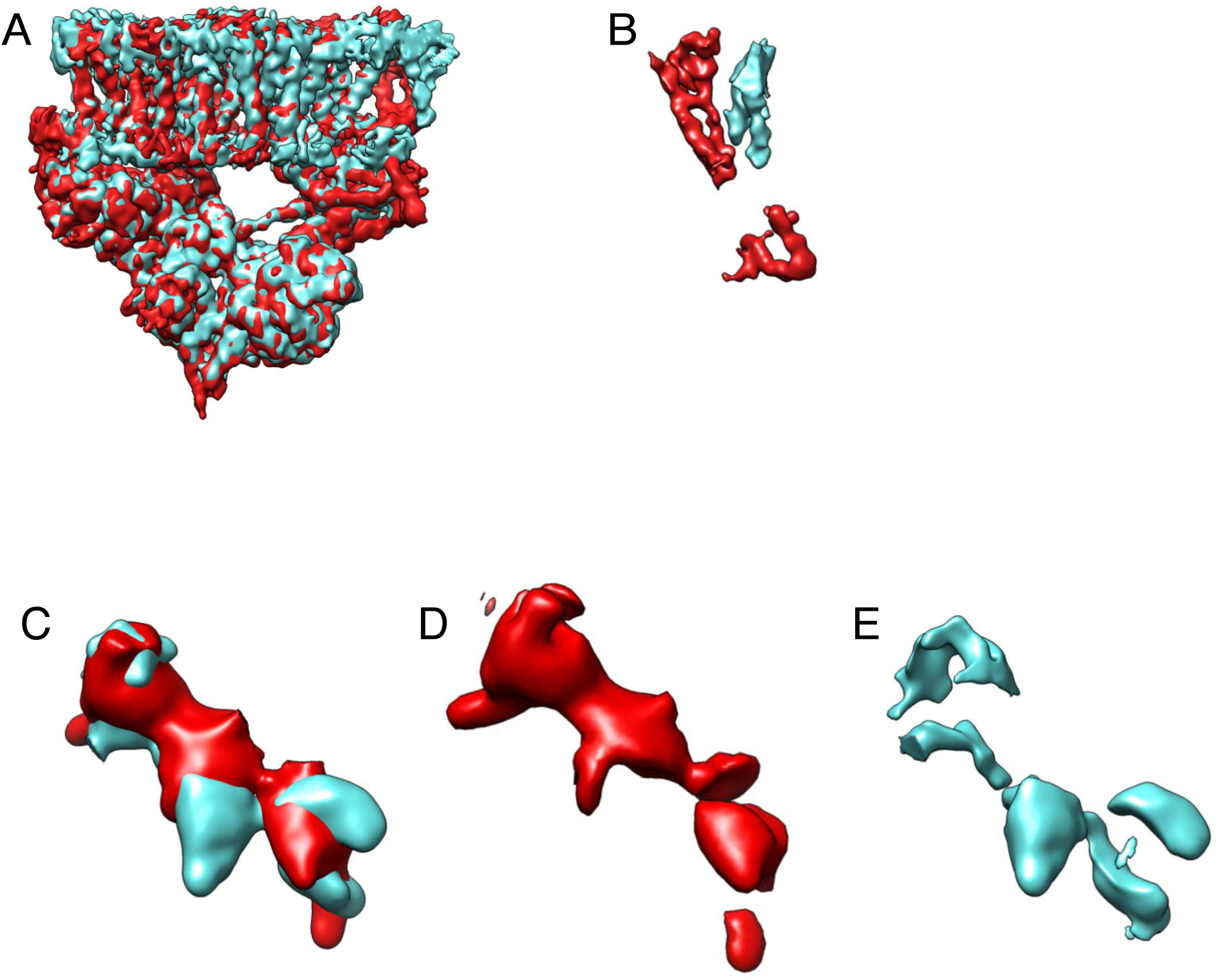
Conformational differences between protomer i and protomer ii. (A) Overlay of the focused refined maps of the transmembrane and upper cytoplasmic regions of protomer i (red) and protomer ii (blue). The two protomers are nearly identical except in the transmembrane domains of EccC_3_ and the N-terminal tail of EccB_3_. (B) Density subtraction of the two protomers in A highlights the regions of difference. (C) Overlay of the ATPase 1,2, and 3 domains of EccC3 from protomer i (red) and protomer ii (blue). (D) Density subtraction of protomer ii from protomer i (E) Density subtraction of protomer i from protomer ii. The lack of overlap between the ATPase domains from the two protomers suggests a different conformation.

**Fig. S11.**
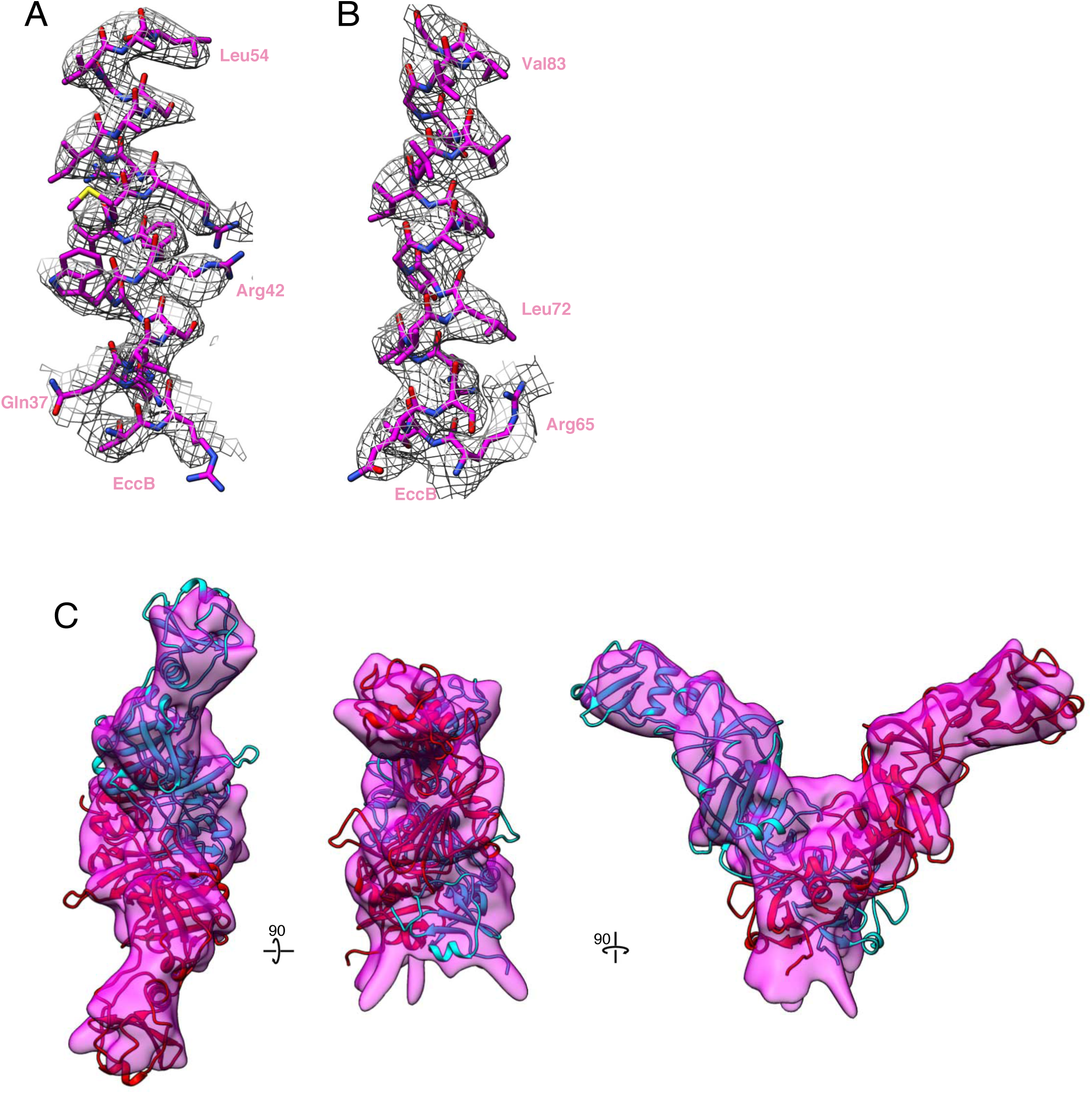
EccB_3_ maps and models. (A) Fit between map and model in the linker α-helix, amino acids 34 to 54. (B) Fit between map and model in the transmembrane helix, amino acids 65 to 85. (C) Fit between homology models of the soluble domain of EccB_3_ and the periplasmic focused refinement map.

**Table S1.** Data collection and refinement statistics. Collection parameters for initial test data set on the Talos Arctica and final collection on Titan Krios microscopes. Refinement details for initial model, consensus map, and focused refinements

| Data Collection |  |  |  |
| --- | --- | --- | --- |
| Collection | Initial Screening | Collection 1 | Collection 2 |
| Microscope | Talos Arctica | Titan Krios | Titan Krios |
| Voltage (kV) | 200 | 300 | 300 |
| Detector | Gatan K2 | Gatan K2 | Gatan K2 |
| Pixel size (Å/pixel) | 1.14 | 0.82 | 0.82 |
| Exposure Time (s) | 9 | 10 | 10 |
| Electron dose (e-/Å<sup>2</sup>) | 63 | 80 | 67 |
| Defocus range (µm) | 1.5-2.5 | 0.4-1.2 | 0.6-1.4 |
| Number of micrographs | 2499 | 2705 | 4632 |
| Consensus Reconstruction |  |  |  |
| Data Set | Initial Screening | Collection 1 & 2 |  |
| Software | Relion 2.1, cisTEM, and cryosparc | Relion 3.0 |  |
| # of particles, picked | 240,000 | 778,149 |  |
| # of particles post, Class2D | 138,000 | 554,901 |  |
| # of particles post, Class3D | 46,830 | 362,438 |  |
| # of particles post, skip align Class3D | NA | 90,479 |  |
| Symmetry | C1 | C1 |  |
| Map sharpening B-factor (Å<sup>2</sup>) | NA | -160 |  |
| Final resolution (Å) | 4.7 Å | 4.0 Å |  |
| Focused Refinements |  |  |  |
| Location of focus | # of particles | Reconstruction resolution (Å) |  |
| Protomer i | 76,967 | 3.8 |  |
| Protomer ii | 66,239 | 3.7 |  |
| Symmetry expanded protomer | 52,067 | 3.7 |  |
| Periplasmic EccB | 70,000 | 5.5 |  |
| ATPase 1,2, and 3 | 30,000 | 10 |  |

**Table S2.** Model Refinement Statistics. Final refinement validation for all amino acids which were modeled de novo (i.e. the translocon pore and translocon gate).

| **Model Refinement** |  |
| --- | --- |
| Total amino acids | 3,156 |
| All-atom clashscore | 13.49 |
| Ramachandran outliers | 0.00% |
| Ramachandran allowed | 5.39% |
| Ramachandran favored | 94.61% |
| Rotamer Outliers | 0.00% |
| Cbeta Deviations | 0% |
| Cis-proline | 0% |
| Cis-general | 0.00% |
| Twisted proline | 0% |
| Twisted general | 0.00% |

**Table S3.**
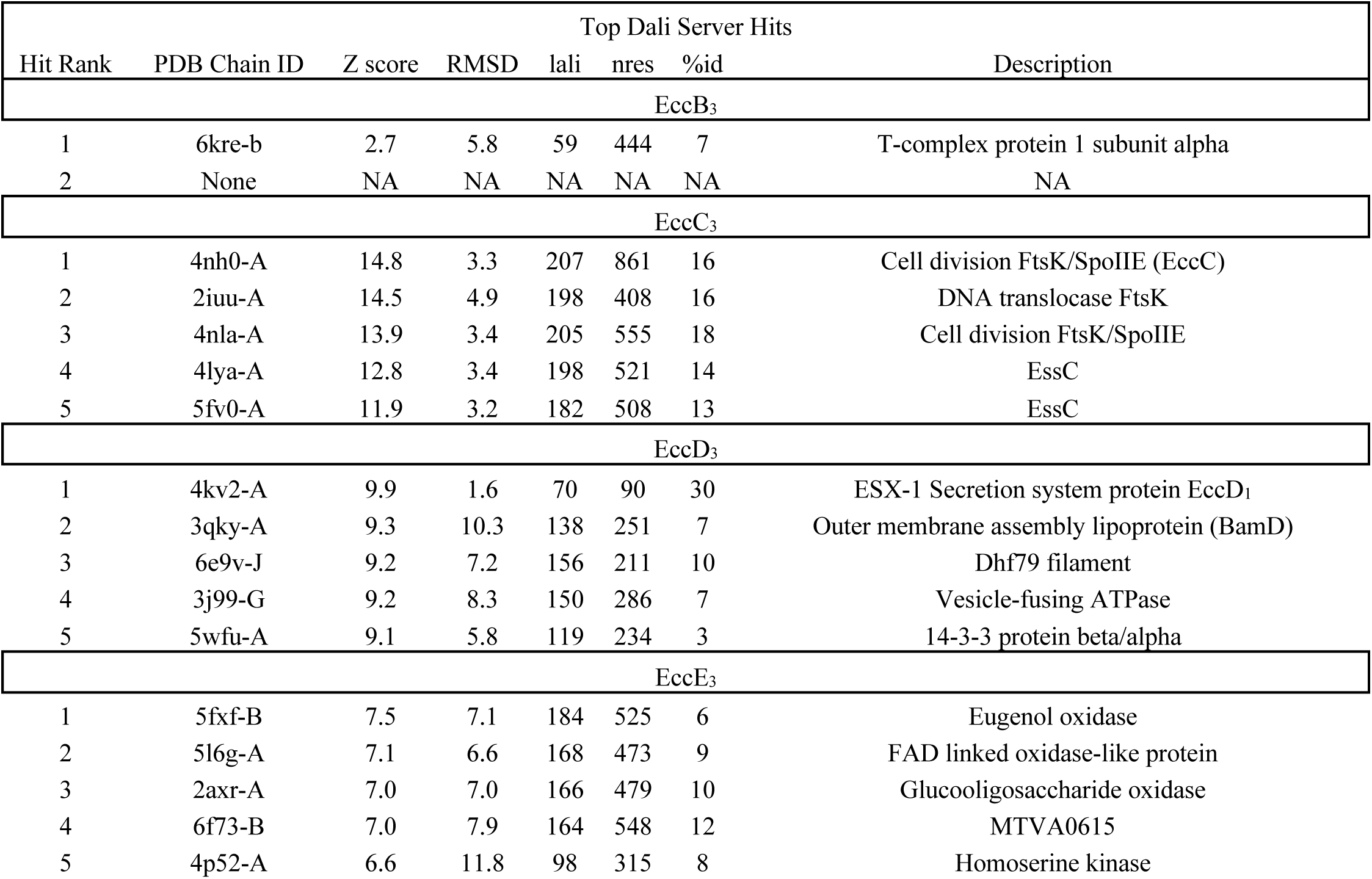
Top Dali server hits. Top 5 non-redundant hits from the Dali server compared against the full PDB for the *de novo* built sections of EccB_3_, EccC_3_, EccD_3_, and EccE_3_.

**Table S4.**
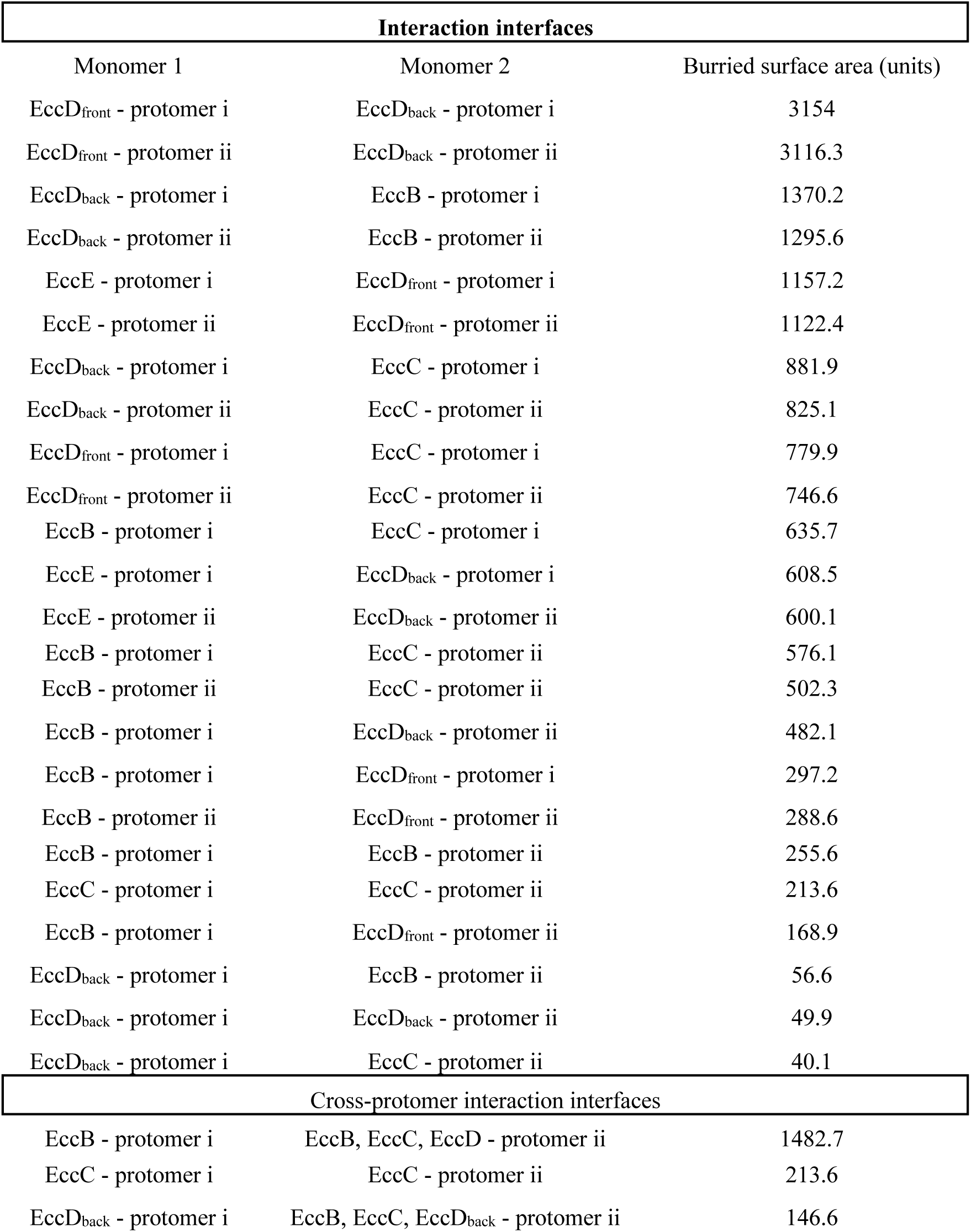
Buried surface area. Buried surface area between subunits as measured by PISA.

## Bibliography

1. M. I. Gröschel, F. Sayes, R. Simeone, L. Majlessi, R. Brosch, ESX secretion systems: mycobacterial evolution to counter host immunity. Nat. Rev. Microbiol. 14, 677–691 (2016).

2. S. A. Stanley, S. Raghavan, W. W. Hwang, J. S. Cox, Acute infection and macrophage subversion by Mycobacterium tuberculosis require a specialized secretion system. Proc. Natl. Acad. Sci. U. S. A. 100, 13001–13006 (2003).

3. K. N. Lewis et al., Deletion of RD1 from Mycobacterium tuberculosis mimics bacille Calmette-Guérin attenuation. J. Infect. Dis. 187, 117–123 (2003).

4. K. M. Guinn et al., Individual RD1-region genes are required for export of ESAT-6/CFP-10 and for virulence of Mycobacterium tuberculosis. Mol. Microbiol. 51, 359–370 (2004).

5. T. Hsu et al., The primary mechanism of attenuation of bacillus Calmette-Guerin is a loss of secreted lytic function required for invasion of lung interstitial tissue. Proc. Natl. Acad. Sci. USA. 100, 12420–12425 (2003).

6. D. Bottai, M. I. Gröschel, R. Brosch, Type VII Secretion Systems in Gram-Positive Bacteria. Curr. Top. Microbiol. Immunol. 404, 235–265 (2017).

7. W. Bitter et al., Systematic genetic nomenclature for type VII secretion systems. PLoS Pathog. 5, e1000507 (2009).

8. E. N. G. Houben et al., Composition of the type VII secretion system membrane complex. Mol. Microbiol. 86, 472–484 (2012).

9. C. Poulsen, S. Panjikar, S. J. Holton, M. Wilmanns, Y.-H. Song, WXG100 protein superfamily consists of three subfamilies and exhibits an α-helical C-terminal conserved residue pattern. PLoS One. 9, e89313 (2014).

10. S. M. Fortune et al., Mutually dependent secretion of proteins required for mycobacterial virulence. Proc. Natl. Acad. Sci. USA. 102, 10676–10681 (2005).

11. T. A. Sysoeva, M. A. Zepeda-Rivera, L. A. Huppert, B. M. Burton, Dimer recognition and secretion by the ESX secretion system in Bacillus subtilis. Proc. Natl. Acad. Sci. USA. 111, 7653–7658 (2014).

12. X.-L. Zhang et al., Core component EccB1 of the Mycobacterium tuberculosis type VII secretion system is a periplasmic ATPase. FASEB J. 29, 4804–4814 (2015).

13. O. S. Rosenberg et al., Substrates Control Multimerization and Activation of the Multi-Domain ATPase Motor of Type VII Secretion. Cell. 161, 501–512 (2015).

14. M. Zoltner et al., EssC: domain structures inform on the elusive translocation channel in the Type VII secretion system. Biochem. J. 473, 1941–1952 (2016).

15. J. M. Wagner et al., Structures of EccB1 and EccD1 from the core complex of the mycobacterial ESX-1 type VII secretion system. BMC Struct Biol. 16, 5 (2016).

16. K. S. H. Beckham et al., Structure of the mycobacterial ESX-5 type VII secretion system membrane complex by single-particle analysis. Nat. Microbiol. 2, 17047 (2017).

17. D. Ilghari et al., Solution structure of the Mycobacterium tuberculosis EsxG·EsxH complex: functional implications and comparisons with other M. tuberculosis Esx family complexes. J. Biol. Chem. 286, 29993–30002 (2011).

18. D. C. Ekiert, J. S. Cox, Structure of a PE-PPE-EspG complex from Mycobacterium tuberculosis reveals molecular specificity of ESX protein secretion. Proc. Natl. Acad. Sci. USA. 111, 14758–14763 (2014).

19. M. Solomonson et al., Structure of the mycosin-1 protease from the mycobacterial ESX-1 protein type VII secretion system. J. Biol. Chem. 288, 17782–17790 (2013).

20. M. S. Siegrist et al., Mycobacterial Esx-3 is required for mycobactin-mediated iron acquisition. Proc. Natl. Acad. Sci. USA. 106, 18792–18797 (2009).

21. A. Serafini, D. Pisu, G. Palù, G. M. Rodriguez, R. Manganelli, The ESX-3 secretion system is necessary for iron and zinc homeostasis in Mycobacterium tuberculosis. PLoS One. 8, e78351 (2013).

22. E. Tinaztepe et al., Role of Metal-Dependent Regulation of ESX-3 Secretion in Intracellular Survival of Mycobacterium tuberculosis. Infect. Immun. 84, 2255–2263 (2016).

23. J. M. Tufariello et al., Separable roles for Mycobacterium tuberculosis ESX-3 effectors in iron acquisition and virulence. Proc. Natl. Acad. Sci. USA. 113, E348–57 (2016).

24. X. Li et al., Transcriptome Landscape of Mycobacterium smegmatis. Front. Microbiol. 8, 2505 (2017).

25. G. M. Rodriguez, M. I. Voskuil, B. Gold, G. K. Schoolnik, I. Smith, ideR, An essential gene in mycobacterium tuberculosis: role of IdeR in iron-dependent gene expression, iron metabolism, and oxidative stress response. Infect. Immun. 70, 3371–3381 (2002).

26. R. Pandey, G. M. Rodriguez, IdeR is required for iron homeostasis and virulence in Mycobacterium tuberculosis. Mol. Microbiol. 91, 98–109 (2014).

27. D. Bottai, A. Serafini, A. Cascioferro, R. Brosch, R. Manganelli, Targeting type VII/ESX secretion systems for development of novel antimycobacterial drugs. Curr. Pharm. Des. 20, 4346–4356 (2014).

28. O. Dussurget, M. Rodriguez, I. Smith, An ideR mutant of Mycobacterium smegmatis has derepressed siderophore production and an altered oxidative-stress response. Mol. Microbiol. 22, 535–544 (1996).

29. K. C. Murphy et al., ORBIT: a new paradigm for genetic engineering of mycobacterial chromosomes. MBio. 9 (2018), doi:10.1128/mBio.01467-18.

30. P. A. Eyers, J. M. Murphy, The evolving world of pseudoenzymes: proteins, prejudice and zombies. BMC Biol. 14, 98 (2016).

31. V. J. C. van Winden et al., Mycosins Are Required for the Stabilization of the ESX-1 and ESX-5 Type VII Secretion Membrane Complexes. MBio. 7 (2016), doi:10.1128/mBio.01471-16.

## References

1. S. Q. Zheng et al., MotionCor2: anisotropic correction of beam-induced motion for improved cryo-electron microscopy. Nature Methods 14, 331 (2017).

2. A. Rohou, N. Grigorieff, CTFFIND4: Fast and accurate defocus estimation from electron micrographs. J Struct Biol 192, 216–221 (2015).

3. J. Zivanov et al., New tools for automated high-resolution cryo-EM structure determination in RELION-3. eLife 7, e42166 (2018).

4. T. Grant, A. Rohou, N. Grigorieff, cisTEM, user-friendly software for single-particle image processing. eLife 7, e35383 (2018).

5. A. Punjani, J. L. Rubinstein, D. J. Fleet, M. A. Brubaker, cryoSPARC: algorithms for rapid unsupervised cryo-EM structure determination. Nature Methods 14, 290 (2017).

6. E. F. Pettersen et al., UCSF Chimera—A visualization system for exploratory research and analysis. Journal of Computational Chemistry 25, 1605–1612 (2004).

7. P. Emsley, B. Lohkamp, W. G. Scott, K. Cowtan, Features and development of Coot. Acta Crystallographica Section D 66, 486–501 (2010).

8. M. Källberg et al., Template-based protein structure modeling using the RaptorX web server. Nature Protocols 7, 1511 (2012).

9. P. V. Afonine et al., Real-space refinement in PHENIX for cryo-EM and crystallography. Acta Crystallographica Section D 74, 531–544 (2018).

10. L. G. Trabuco, E. Villa, E. Schreiner, C. B. Harrison, K. Schulten, Molecular dynamics flexible fitting: A practical guide to combine cryo-electron microscopy and X-ray crystallography. Methods 49, 174–180 (2009).

11. R. T. Kidmose et al., Namdinator - Automatic Molecular Dynamics flexible fitting of structural models into cryo-EM and crystallography experimental maps. bioRxiv, 501197 (2018).

12. L. Pravda et al., MOLEonline: a web-based tool for analyzing channels, tunnels and pores (2018 update). Nucleic Acids Research 46, W368–W373 (2018).

13. E. Krissinel, Stock-based detection of protein oligomeric states in jsPISA. Nucleic Acids Research 43, W314–W319 (2015).

14. L. Holm, Benchmarking fold detection by DaliLite v.5. Bioinformatics, (2019).

15. T. D. Goddard et al., UCSF ChimeraX: Meeting modern challenges in visualization and analysis. Protein Science 27, 14–25 (2018).

